# Effects of dual deletion of *glnR* and *mtrA* on expression of nitrogen metabolism genes in *Streptomyces venezuelae*

**DOI:** 10.1101/2021.10.15.464524

**Authors:** Yanping Zhu, Jiao Wang, Wenya Su, Ting Lu, Aiying Li, Xiuhua Pang

**Affiliations:** The State Key Laboratory of Microbial Technology, Shandong University, Qingdao 266237, China

**Keywords:** *Streptomyces*, nitrogen metabolism, MtrA, GlnR

## Abstract

GlnR activates nitrogen metabolism genes under nitrogen-limited conditions whereas MtrA represses these genes under nutrient-rich conditions in *Streptomyces*. In this study, we compared the transcription patterns of nitrogen metabolism genes in a double deletion mutant (Δ*mtrA*-*glnR*) lacking both *mtrA* and *glnR* and in mutants lacking either *mtrA* (Δ*mtrA*) or *glnR* (Δ*glnR*). The nitrogen metabolism genes were expressed similarly in Δ*mtrA*-*glnR* and Δ*glnR* under both nitrogen-limited and nutrient-rich conditions, with patterns distinctly different from that of Δ*mtrA*, suggesting a decisive role for GlnR in the control of nitrogen metabolism genes and further suggesting that regulation of these genes by MtrA is GlnR-dependent. MtrA and GlnR utilize the same binding sites upstream of nitrogen metabolism genes, and we showed stronger *in vivo* binding of MtrA to these sites under nutrient-rich conditions and of GlnR under nitrogen-limited conditions, consistent with the higher levels of MtrA or GlnR under those respective conditions. In addition, we showed that both *mtrA* and *glnR* are auto-regulatory. Our study provides new insights into the regulation of nitrogen metabolism genes in *Streptomyces*.

## Introduction

Nitrogen sources, whether organic or inorganic, are a necessity for all living organisms, including microbes, and the regulation of nitrogen metabolism is complex and varied in bacteria (Leigh & Dodsworth, 2007, Merrick & Edwards, 1995). Under nitrogen-limited growth conditions, genes involved in nitrogen assimilation are expressed to enable the acquisition and conversion of inorganic nitrogen sources into organic nitrogen sources such as glutamine and glutamate.*Streptomyces* are a genus of Gram-positive and filamentous actinobacteria mostly known for their potential in producing antibiotics as well as for their complex development cycle, including spore formation (Chater, 2011, Hopwood, 2007). Within members of this genus, multiple nitrogen assimilation genes have been identified, including *amtB*, encoding a protein that transports extracellular ammonium into the cell; *narGHIJ*, encoding nitrate reductase, which reduces nitrate into nitrite; *nirBCD*, encoding nitrite reductase, which reduces nitrite into ammonium; *ureABC*,encoding a urease for the cleavage of urea into NH_4_^+^; *glnA* and *glnII*, both encoding a glutamine synthetase, and *gltDB*, encoding a glutamate synthase, which synthesize glutamine or glutamate, respectively, using NH_4_^+^ imported from the extracellular environment or converted from nitrate,nitrile, or urea (Wolfgang Wohlleben, 2011).

Under nitrogen-limited conditions, most *Streptomyces* genes for nitrogen assimilation are activated by the orphan response regulator GlnR (Wolfgang Wohlleben, 2011, Tiffert *et al*., 2008,Fink *et al*., 2002). GlnR boxes were identified for *amtB, glnII*, and other nitrogen assimilation genes, indicating that these nitrogen metabolism genes are targeted by GlnR (Tiffert *et al*., 2008, Fink *et al*., 2002, Pullan *et al*., 2011, Tiffert *et al*., 2011). Under nutrient-rich growth conditions, surplus nitrogen is present, and therefore nitrogen assimilation genes do not need to be expressed and appear to be silent. We revealed that this silencing, or only basal level of expression of nitrogen assimilation genes, is the result of repression by MtrA (Zhu *et al*., 2019), a global response regulator that is also required for cellular development (Zhang *et al*., 2017), antibiotic production (Zhu *et al*., 2020a, Som *et al*., 2017a, Som *et al*., 2017b), and phosphate metabolism (Zhu *et al*., 2021). Intriguingly, the sequence recognized by MtrA (MtrA site) is similar to the GlnR box (Zhang *et al*., 2017), and thus MtrA can interact with the GlnR boxes upstream of the nitrogen metabolism genes that are targeted by GlnR (Zhu *et al*., 2019), suggesting that MtrA potentially competes with GlnR in the regulation of nitrogen metabolism genes. GlnR and MtrA have been characterized as the two major regulators for nitrogen metabolism in *Streptomyces* and potentially in other actinobacteria (Zhu *et al*., 2019, Wang *et al*., 2015). Studies suggest that these two regulators function under contrasting nitrogen supply conditions, although minor regulatory effects on nitrogen metabolism genes by PhoP and AfsQ1 were also observed under specific conditions (Rodriguez-Garcia *et al*., 2009, Wang *et al*., 2013).

Although it is known that MtrA and GlnR function by binding their target sites under nitrogen-limited and nutrient-rich conditions, respectively, it is not known whether MtrA or GlnR still have a role under the contrasting condition that does not favor their function. The combined regulatory effect of MtrA and GlnR on nitrogen metabolism genes is also unknown. In this study, we investigated the binding of MtrA and GlnR under different nitrogen conditions and explored the combined effect of MtrA and GlnR on nitrogen metabolism genes, thus providing new insights into the understanding of nitrogen metabolism in *Streptomyces*.

## Results

### The role of MtrA on nitrogen metabolism is similar in S. venezuelae and S. coelicolor

Our previous study showed that MtrA represses nitrogen metabolism genes such as *amtB* and *glnII* in *S. coelicolor* and *S. lividans* (Zhu *et al*., 2019), which are closely related species (Kawamoto & Ochi, 1998, Lewis *et al*., 2010). To investigate whether MtrA has a similar regulatory effect in *S. venezuelae*, which is more distantly related to the model strain *S. coelicolor*, we compared the expression levels of known nitrogen metabolism genes in the wild-type strain *S. venezuelae* ATCC10712 and Δ*mtrA*_*SVE*_, which is an *mtrA* deletion mutant of this strain (Zhu *et al*., 2020a). Our transcriptional analysis showed that the nitrogen metabolism genes *amtB, glnK, glnD, nirB*, and *glnII* were more highly expressed in the mutant on rich medium, including YBP (Fig. S1) and R2YE (Fig. 1A). We also investigated whether MtrA recognizes the GlnR boxes of nitrogen metabolism genes in *S. venezuelae*. We showed that MtrA binds the GlnR boxes of nitrogen metabolism genes including *amtB, glnII, glnA, glnR, nirB, ureA, gltB* using the wild-type and mutagenized sequence as probes (Fig. S2-S9), indicating that MtrA recognizes GlnR box and represses these nitrogen metabolism genes in *S. venezuelae*, consistent with its role in *S. coelicolor* (Zhu *et al*., 2019)

**Figure 1.**
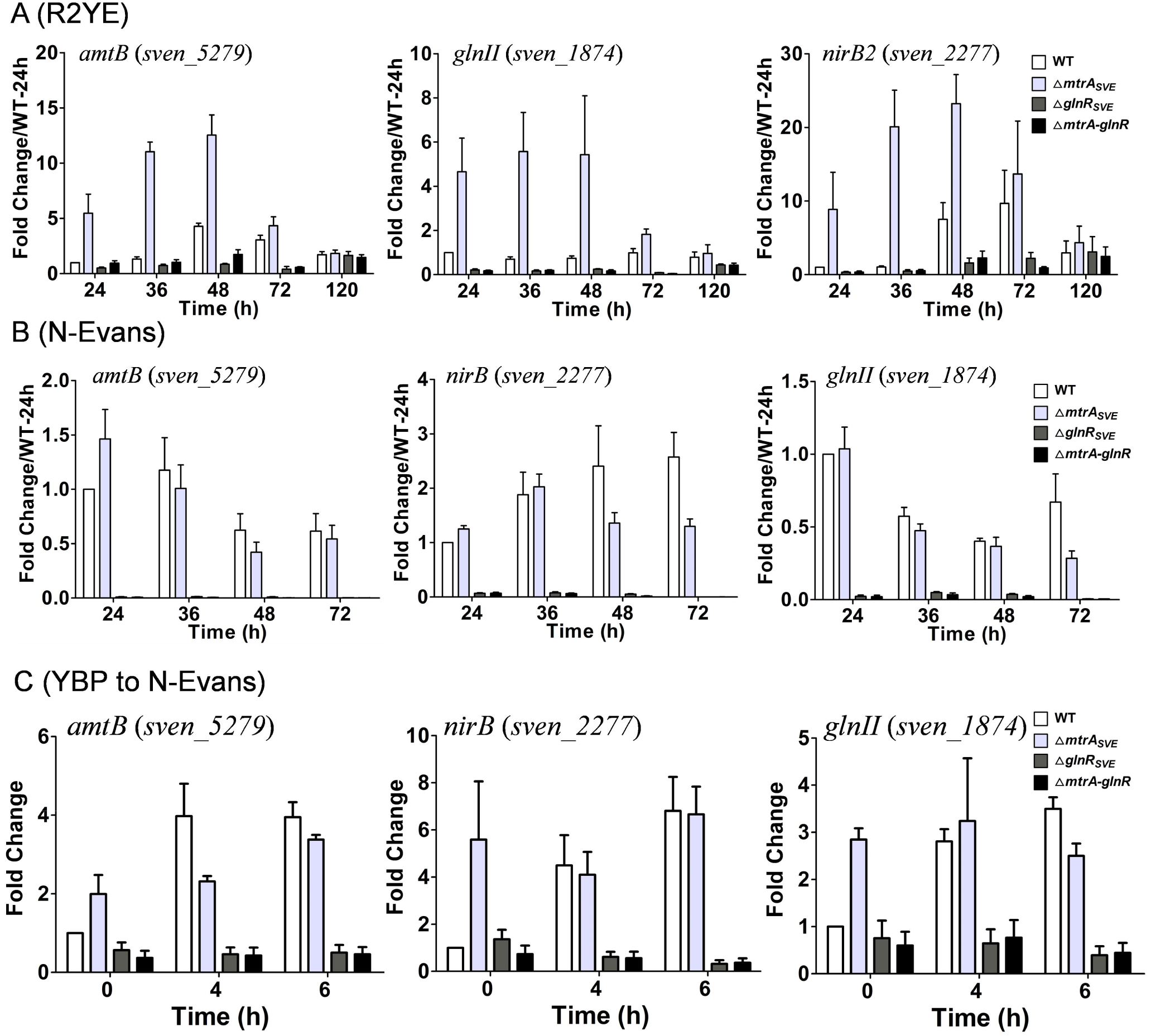
Transcriptional analysis of nitrogen metabolism genes in Δ*mtrA*_*SVE*_, Δ*glnR*_*SVE*_, and Δ*mtrA*-*glnR* mutants by real-time PCR. *Streptomyces* strains were cultured on solid (A) R2YE, (B) N-Evans, or (C) shifted from liquid YBP to liquid N-Evans, and RNA samples from 10712 (WT), Δ*mtrA*_*SVE*_, Δ*glnR*_*SVE*_, and Δ*mtrA*-*glnR* were isolated at the indicated times. Expression of *hrdB*, encoding the major sigma factor, was used as an internal control. For each gene, the expression level in the wild-type strain (WT) at the first time point was arbitrarily set to one. The y-axis shows the fold change in expression in WT, Δ*mtrA*_*SVE*_, Δ*glnR*_*SVE*_, and Δ*mtrA*-*glnR* over the levels in WT at the first time point. Results are the means (±SD) of triplet biological experiments.

### The two nitrogen metabolism regulators mtrA and glnR are auto-regulatory

MtrA functions as a repressor for nitrogen metabolism genes; however, the impact of MtrA on its own expression was not known. To determine whether *mtrA* is auto-regulatory, a set of primers was designed to target a region of *mtrA* still present in Δ*mtrA*_*SVE*_, and these primers were used in transcriptional analysis by real-time PCR. To facilitate comparison, the expression level of *mtrA* at the first time point in the wild-type strain was arbitrarily set to one (Fig. 2). For the wild-type strain, an expression level of about one (ranging from 0.74 to 1.1) was detected for *mtrA* throughout the entire time course, whereas the fold change in expression of *mtrA* in Δ*mtrA*_*SVE*_ ranged from 170±37 to 287±153 on R2YE (Fig. 2A). *mtrA* also maintained a basal expression level throughout the time course for the wild-type strain on the nitrogen-limited medium N-Evans (Fig. 2B), and although its upregulation in Δ*mtrA*_*SVE*_ was not so striking as on R2YE, *mtrA* expression in the mutant ranged from 59±24 to 82±25, indicating that MtrA represses its own expression under both nitrogen-limited and nutrient-rich conditions. Notably, no detectable level of expression was detected for *mtrA* in transcriptional analysis using a second set of primers that target the deleted sequence of *mtrA* (Fig. S10), confirming the removal of a portion of *mtrA*. However, analysis using each set of primers showed that *mtrA* was overexpressed in a *glnR* mutant strain of *S. venezuelae* (Δ*glnR*_*SVE*_) on N-Evans medium (Fig. 2B, Fig. S10A), indicating that GlnR represses *mtrA* under nitrogen-limited conditions, which is in agreement with our previous report on *S. coelicolor* (Zhu *et al*., 2019).

**Figure 2.**
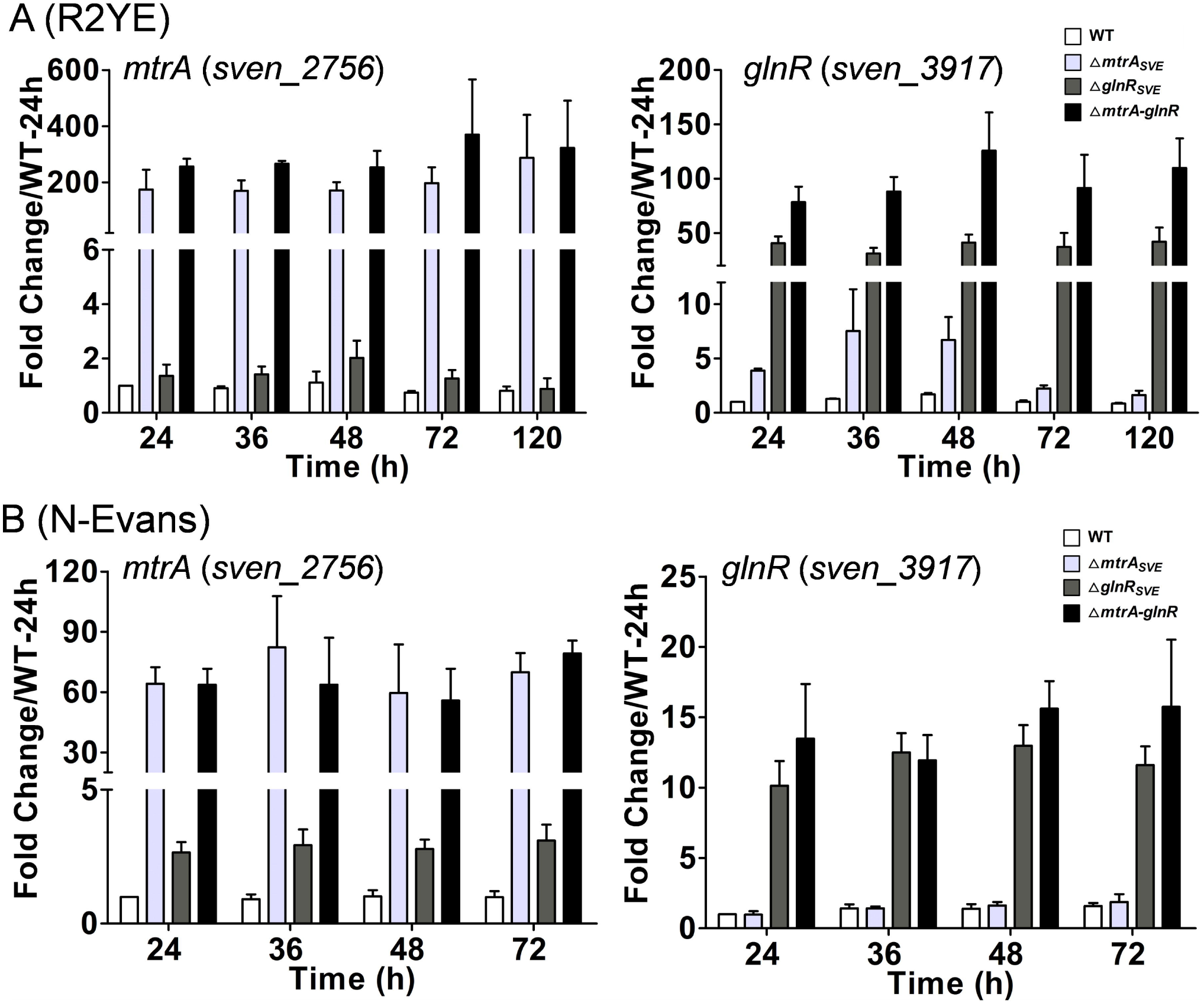
Transcriptional analysis of *mtrA* and *glnR* in Δ*mtrA*_*SVE*_, Δ*glnR*_*SVE*_, and Δ*mtrA*-*glnR* mutants by real-time PCR. *Streptomyces* strains were cultured on solid R2YE (A) and N-Evans (B), and RNA samples from the wild-type strain (WT), Δ*mtrA*_*SVE*_, Δ*glnR*_*SVE*_, and Δ*mtrA*-*glnR* were isolated at the indicated times. Expression of *hrdB*, encoding the major sigma factor, was used as an internal control. For each gene, the expression level in WT at the first time point was arbitrarily set to one. The y-axis shows the fold change in expression in WT, Δ*mtrA*_*SVE*_, Δ*glnR*_*SVE*_, and Δ*mtrA*-*glnR* over the levels in WT at the first time point. Primer sets that target remaining regions of *mtrA* or *glnR* in Δ*mtrA*_*SVE*_ and Δ*glnR*_*SVE*_, respectively, were used. Results are the means (±SD) of triplet biological experiments.

We also investigated whether GlnR is auto-regulatory using a set of primers that targeted at the undeleted sequence of *glnR* and a *glnR* mutant strain of *S. venezuelae* (Zhu *et al*., 2020a). While a level of about one was detected for *glnR* in the wild-type strain through time course on both media, *glnR* reached a level ranging from 10±1.7 to 12.9±1.5 on N-Evans and a level ranging from 31±5.2 to 42±12.9 on R2YE in Δ*glnR*_*SVE*_ (Fig. 2), indicating that GlnR represses its own expression under both nitrogen-limited and nutrient-rich conditions. However, when a second set of primers that targets a segment of *glnR* deleted in Δ*glnR*_*SVE*_ was used, no detectable level of expression was detected for *glnR* (Fig. S10), confirming the deletion. Overexpression of *glnR* was detected in the *mtrA* mutant strain using both sets of primers on R2YE (Fig. 2A, Fig. S10B), validating that MtrA represses *glnR* in *S. venezuelae* under nutrient-rich conditions, similar to our findings with *S. coelicolor* (Zhu *et al*., 2019).

### mtrA is the target of both MtrA and GlnR in S. venezuelae

To explore the potential mechanism for the autoregulation of *mtrA*, the sequence upstream of *mtrA* was examined, and a potential MtrA site was identified (Fig. 3A). To determine if MtrA interacts with this site, an electrophoretic mobility shift assay (EMSA) was performed using purified MtrA and short oligonucleotides containing the predicted MtrA site as probe. The EMSA analysis showed that MtrA binds the probe with the predicted MtrA site, but not the probes with mutations at the conserved nucleotides of the site, upstream of *mtrA in vitro* (Fig. 3B). To determine whether MtrA also binds this site *in vivo*, the integrative plasmid pMtrA-FLAG, which expresses an MtrA-FLAG fusion protein under the control of the native *mtrA* promoter, was constructed and was introduced into Δ*mtrA*_*SVE*_. Whereas Δ*mtrA*_*SVE*_ exhibited a delay in spore formation and defective pigment production, these defects were reversed in MtrA-FLAG-Δ*mtrA*_*SVE*_ (Fig. S11), indicating that FLAG-tagged MtrA is expressed and functional. MtrA-FLAG-Δ*mtrA*_*SVE*_ was therefore used for ChIP analysis with anti-FLAG antibody. The binding level detected at the *mtrA* promoter remained at around background levels for the control wild-type strain, whereas a relative binding level of about four was detected for MtrA-FLAG-Δ*mtrA*_*SVE*_ on R2YE and N-Evans (Fig. 4A), indicating that MtrA binds this site *in vivo* under both conditions.

**Figure 3.**
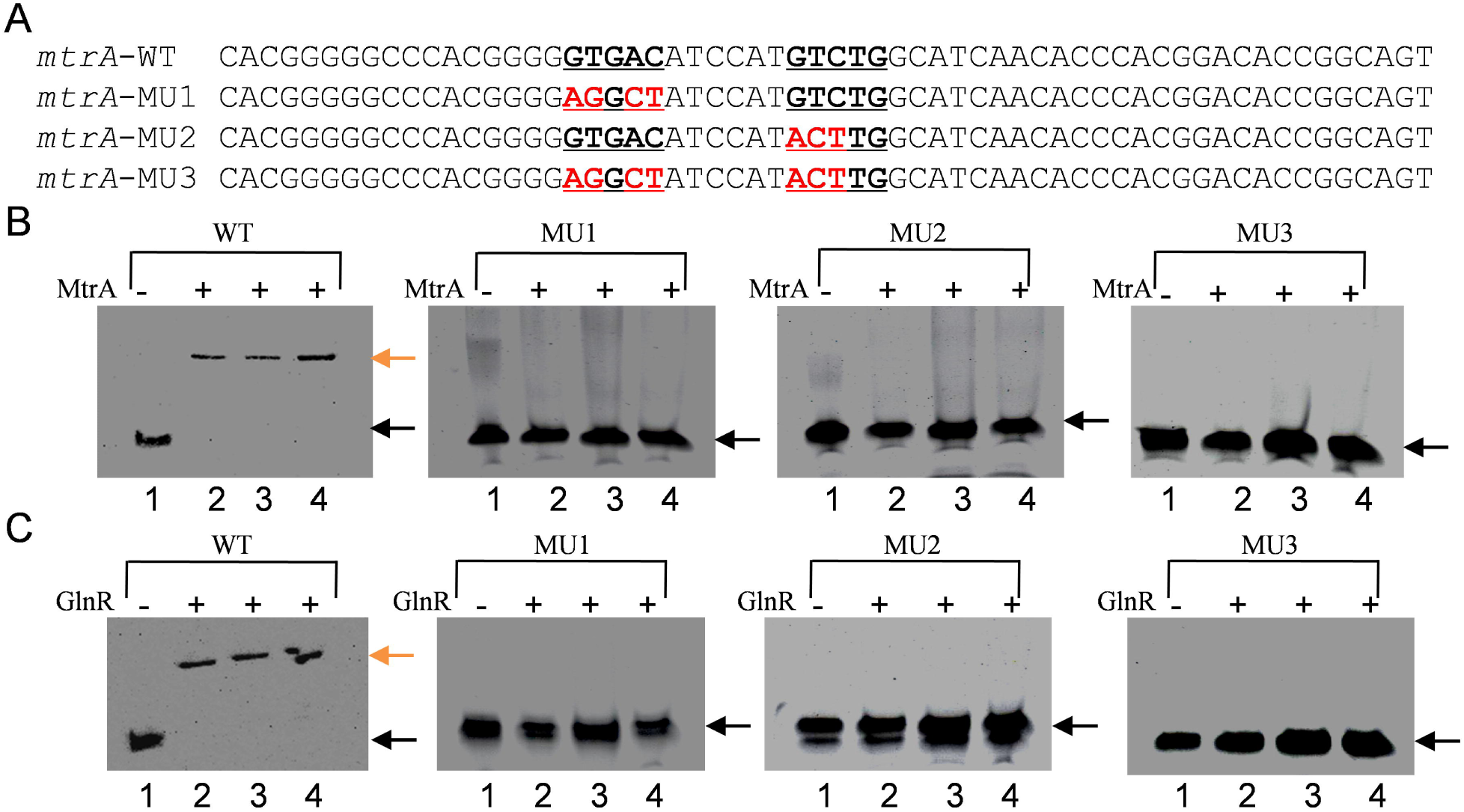
*mtrA* is a target of MtrA and GlnR. (A) the predicted MtrA site upstream of *mtrA* in *S. venezuelae*. The MtrA site is underlined and in boldface, and tested mutations are highlighted in red. (B) EMSA with MtrA and 59-bp probes containing the predicted MtrA site or the MtrA site with mutations. Reactions were carried out with the addition of no MtrA (lane 1), or with 1.18 μM (lane 2), 4.74 μM (lane 3), or 8.29 μM (lane 4) MtrA. (C) EMSA with GlnR and a 59-bp probe containing the predicted MtrA site or the MtrA site with mutations. Reactions were carried out with the addition of no GlnR (lane 1), or with 0.45 μM (lane 2), 1.78 μM (lane 3), or 3.11 μM (lane 4) GlnR. The red and black arrows indicate the positions of the shifted and free probes, respectively.

**Figure 4.**
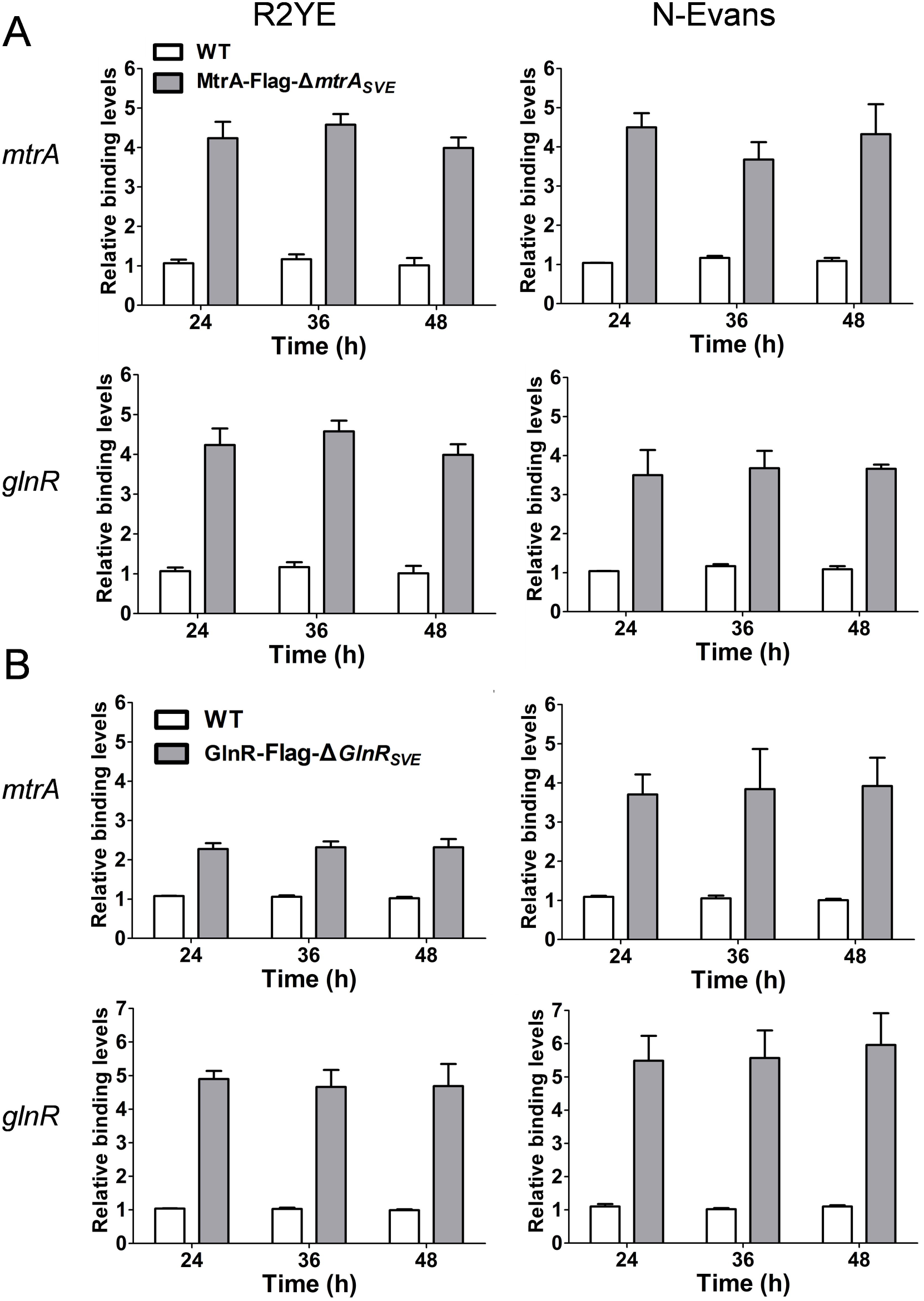
ChIP-qPCR analysis of MtrA and GlnR binding to the *mtrA* and *glnR* promoters. The wild-type strain (WT), MtrA-FLAG-Δ*mtrA*_*SVE*_, and GlnR-FLAG-Δ*glnR*_*SVE*_ were cultured on solid R2YE and N-Evans media and were processed at the indicated times. The y-axis shows the binding level of MtrA or GlnR relative to the background in WT, MtrA-FLAG-Δ*mtrA*_*SVE*_, and GlnR-FLAG-Δ*glnR*_*SVE*_. As WT contains native MtrA and GlnR only, the results for this strain are equivalent to background amplification of the target sequences The data show the means (±SD) of triplet biological experiments.

To determine whether GlnR also binds the MtrA site upstream of *mtrA in vivo*, the integrative plasmid pGlnR-FLAG expressing a GlnR-FLAG fusion protein under the control of the native *glnR* promoter was constructed and introduced into Δ*glnR*_*SVE*_. The delay in spore formation and the defect in pigment production by Δ*glnR*_*SVE*_ were restored to nearly wild-type levels in GlnR-FLAG-Δ*glnR*_*SVE*_ (Fig. S12), indicating that the FLAG-tagged GlnR was expressed and functional. This strain was then used for ChIP analysis. Our data showed that GlnR bound the MtrA site upstream of *mtrA in vitro* (Fig. 3C) and also *in vivo* on both R2YE and N-Evans (Fig. 4B). Altogether, we showed that the *mtrA* promoter is a target of both MtrA and GlnR, explaining the autoregulation of MtrA and its regulation by GlnR.

### Transcription of nitrogen metabolism genes in ΔmtrA-glnR

Previously, we used mutant strains with deletions of either *mtrA* or *glnR* to investigate the impact of MtrA or GlnR, respectively, on nitrogen metabolism genes (Zhu *et al*., 2019). In this study, we asked what is the combined impact of MtrA and GlnR on nitrogen metabolism genes? To address this question, we generated the mutant strain Δ*mtrA*-*glnR*, which has deletions of both *mtrA* and *glnR*. Next, we compared the expression of nitrogen metabolism genes in this strain and in the single *mtrA* and *glnR* deletion mutants. To facilitate the comparison, the expression level of each gene in the wild-type strain at the first time point was arbitrarily set to one. As noted previously, on R2YE, nitrogen genes such as *amtB, glnII*, and *nirB2* were markedly upregulated in Δ*mtrA*_*SVE*_, most notably at the three early time points (24, 36, 48 h) (Fig. 1A), confirming a major role for MtrA under nutrient-rich conditions. In contrast, the expression level of these genes was reduced moderately, mostly at two time points (48 and 72 h) in Δ*glnR*_*SVE*_, suggesting a positive, although minor, role for GlnR in their regulation under nutrient-rich conditions. Unexpectedly, in Δ*mtrA*-*glnR*, the expression levels of several nitrogen metabolism genes on R2YE, including *amtB, nirB*, and *glnII*, were more similar to those in Δ*glnR*_*SVE*_, a transcription pattern distinctly different from that observed in Δ*mtrA*_*SVE*_ (Fig. 1A and Fig. S13). Two exceptions were *gltB* and *gdhA*, whose transcription pattern in Δ*mtrA*-*glnR* was more similar to that in Δ*mtrA*_*SVE*_ at several time points (Fig. S13).

We next examined the transcription patterns in Δ*mtrA*-*glnR* grown on N-Evans, a nitrogen-limited medium on which GlnR functions as an activator for nitrogen metabolism genes (Zhu *et al*., 2019, Tiffert *et al*., 2008). In Δ*mtrA*_*SVE*_, the expression level of *amtB, nirB*, and *glnII* was either comparable to that of the wild-type control or only slightly altered (Fig. 1B and Fig. S14), suggesting a minor role for MtrA under nitrogen-limited conditions. As expected, only a minimal level of expression was detected for these genes in Δ*glnR*_*SVE*_ (Fig. 1B and Fig. S14), consistent with the major role for GlnR under nitrogen-limited conditions. Similar to the results for Δ*glnR*_*SVE*_, only minimal expression of these genes was detected in Δ*mtrA*-*glnR* on N-Evans (Fig. 1B and Fig. S14). The exception was *gltB*, which was upregulated in both Δ*mtrA*_*SVE*_ and Δ*glnR*_*SVE*_ (Fig. S14) but which showed even higher expression in the double mutant at the two early time points (24 h and 36 h), suggesting a synergistic effect from the loss of *mtrA* and *glnR*.

We next investigated the expression profiles of nitrogen metabolism genes in Δ*mtrA*-*glnR* following a shift from nutrient-rich (YBP broth) to nitrogen-limited conditions (N-Evans broth) (Fig. 1C and Fig. S15). RNA extracts were prepared directly from YBP and designated as the time 0 samples, prior to the transfer of culture to N-Evans broth for further growth for four or six hours. The expression level of each gene in the wild-type control at time 0 was arbitrarily set to one. As expected, the expression levels of nitrogen metabolism genes were all increased at time 0 in Δ*mtrA*_*SVE*_, and levels were either comparable to the wild-type strain or only slightly impacted in Δ*glnR*_*SVE*_, with the expression patterns in Δ*mtrA*-*glnR* similar to those in Δ*glnR*_*SVE*_ (Fig. 1C and Fig. S15). After four and six hours of growth in N-Evans, the expression profiles of these genes in Δ*mtrA*-*glnR* remained nearly identical to those of Δ*glnR*_*SVE*_. In conclusion, the transcriptional patterns of nitrogen metabolism genes in Δ*mtrA*-*glnR* is similar to that in Δ*glnR*_*SVE*_ under nitrogen-limited or nutrient-rich conditions, or under a nutrient shift from nutrient-rich to nitrogen-limited conditions, implying a decisive role for GlnR in the regulation of nitrogen metabolism genes.

### The relative expression levels of mtrA, glnR, and nitrogen metabolism genes under different growth conditions

Our previous and this study showed that MtrA plays a major role in nitrogen metabolism under nutrient-rich conditions and a minor role under nitrogen-limited conditions. However, it was not known whether MtrA is differentially expressed under these two conditions. Therefore, the expression level of *mtrA* on different growth media was compared using the wild-type strain. The expression level of *hrdB*, which served as the internal control, was arbitrarily set to one at each time point, and the expression of *mtrA* at each time point was calculated relative to *hrdB* (Fig. 5). On YBP, the expression levels of *mtrA* were higher (0.44-0.54) at the two early time points than at the later time points (0.18-0.28). On R2YE, a level ranging from 0.44-0.49 was detected for *mtrA* at the three early time points, whereas the two later time points were lower (0.27-0.32), suggesting highest expression of *mtrA* at the early growth phase under nutrient-rich conditions. On N-Evans, levels remained around 0.26-0.28 for *mtrA*, which is comparable to the levels at the two later time points on R2YE (Fig. 5A), suggesting that, although *mtrA* plays a minor role under nitrogen-limited conditions, it is still moderately expressed under such conditions and its role is also influenced by growth phase.

**Figure 5.**
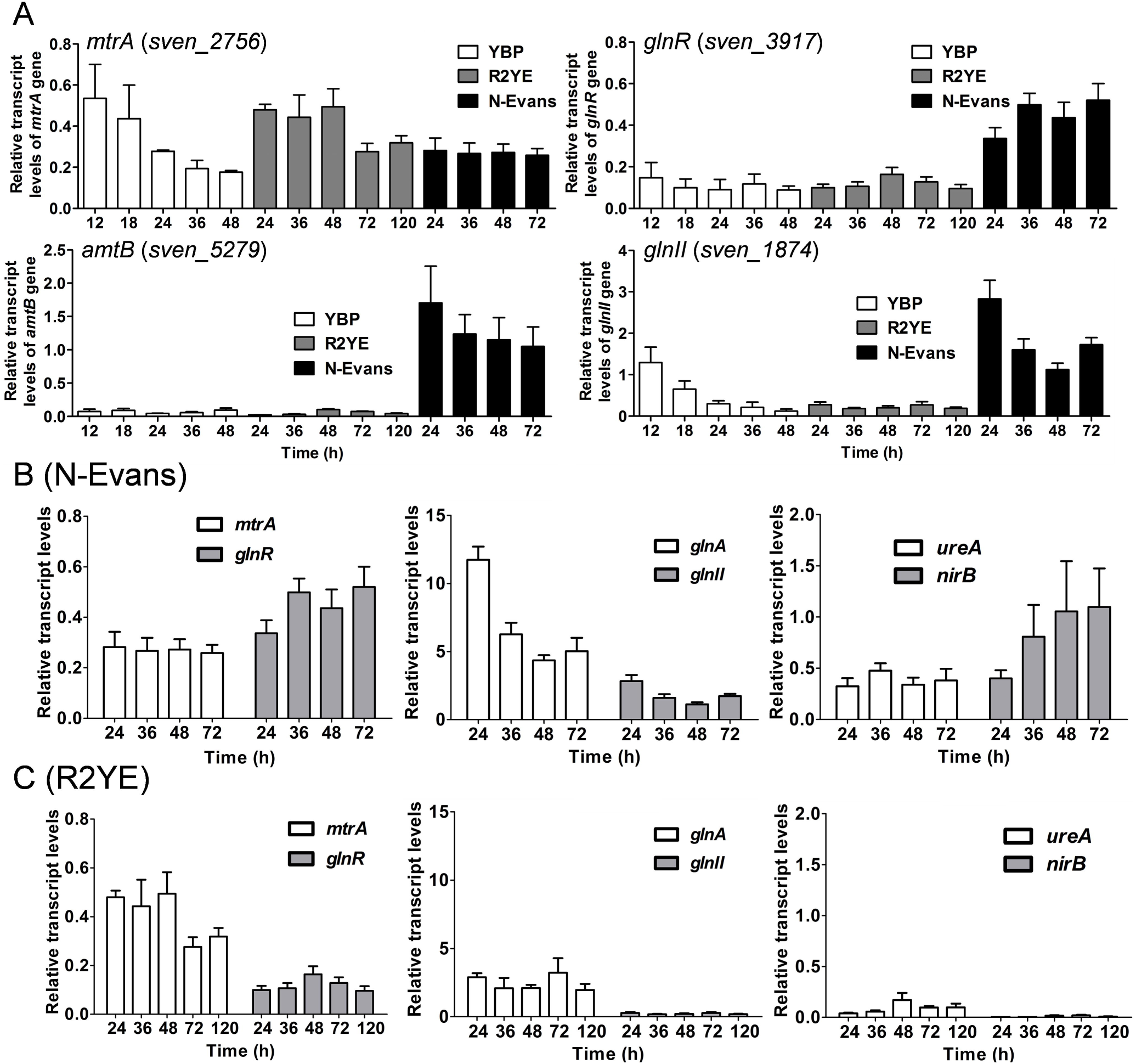
Transcriptional analysis of *mtrA, glnR*, and nitrogen metabolism genes in the wild-type strain 10712 grown on different media. (A) Transcription analysis using YBP, R2YE, and N-Evans cultures. For each gene, the expression level of *hrdB* at each time point was arbitrarily set to one. The y-axis shows the fold change in expression of each gene over the expression level of *hrdB* for each time point. Results are the means (±SD) of triplet biological experiments. (B-C) Transcriptional analysis using R2YE or N-Evans cultures at various time points. Results are the means (±SD) of triplet biological experiments and were calculated as for panel A.

We also examined the response of *glnR* to different nitrogen conditions using the wild-type strain (Fig. 5A). Expression levels of about 0.09-0.14 were detected for *glnR* on YBP medium, which was comparable with its levels on R2YE (0.10-0.16). However, markedly higher levels, ranging from 0.34-0.52, were detected for *glnR* on N-Evans, indicating that, although *glnR* is expressed under nutrient-rich conditions, its expression was much higher in nitrogen-limited conditions. These findings are consistent with a previous report indicating that GlnR is the major regulator for nitrogen metabolism genes under nitrogen-limited conditions but not under nutrient-rich conditions (Tiffert *et al*., 2008). The expression levels of *amtB, glnK, glnD, glnII, nirB*, and *ureA* were also notably higher on N-Evans than on YBP and R2YE (Fig. 5A and Fig. S16), in agreement with a previous report indicating that these nitrogen metabolism genes respond to nitrogen-limited conditions in *S. coelicolor* (Tiffert *et al*., 2008). However, *gdhA* expression was barely detectable on any of the media, while *gltB* expression was detectable and remained at roughly comparable levels on all three media types (Fig. S16), implying that these two genes respond differently from other nitrogen metabolism genes.

The expression levels of *mtrA, glnR*, and other nitrogen metabolism genes under the same growth conditions were also compared (Figs. 5B, 5C, S17, and S18). *mtrA* expression remained lower than *glnR* expression (0.26-0.28 vs 0.33-0.52) at the four time points on N-Evans (Fig. 5B), whereas on R2YE, *mtrA* expression was consistently higher than that of *glnR* (0.27-0.49 vs 0.095-0.164) (Fig. 5C), with the large difference in expression levels indicating that *glnR* is more sensitive to nitrogen availability. Both *glnA* and *glnII* encode glutamine synthetases; however, *glnA* was expressed much more highly than *glnII* on N-Evans (Fig. 5B) and R2YE (Fig. 5C), supporting the notion that GlnA is the major glutamine synthetase in *S. coelicolor* (Tiffert *et al*., 2008).

### MtrA binds differentially to the MtrA/GlnR sites of nitrogen metabolism genes under different growth conditions

To investigate if the differential expression of *mtrA* on different media is reflected at the protein level, we performed Western blot analysis using MtrA-FLAG-Δ*mtrA*_*SVE*_ grown on solid R2YE and N-Evans. Crude cellular lysates were extracted at the same four time points. Whereas the level of MtrA was almost constant on either R2YE or N-Evans, the level on N-Evans was notably lower than on R2YE (Fig. 6A), which is consistent with the transcriptional analysis (Fig. 5). To explore whether the different level of MtrA leads to differential binding *in vivo* to the MtrA/GlnR sites of nitrogen metabolism genes, previously tested in ESMAs (Fig. S2-S9), we performed ChIP-qPCR analysis and compared the binding level of MtrA in cultures grown on R2YE and N-Evans (Fig. 6B, 6C and Fig. S19). Higher binding levels were detected on R2YE than on N-Evans and the levels are fairly constant for a given medium. For example, at 24 h, the levels on R2YE versus N-Evans were as follows: *glnA* (5.02±0.81 vs 3.36±0.83), *glnII* (5.30±0.75 vs 2.58±0.28), *ureA* (4.75±0.67 vs 2.91±0.71), *amtB* (4.56±0.51 vs 3.76±0.64). However, only minor differences in binding by MtrA were observed for *gdhA* and *gltB* (Fig. S19), and no *in vivo* binding was detected for *nirB* and *sven_1860* (Fig. S19), although MtrA bound these two sites *in vitro* (Fig. S6, S9).

**Figure 6.**
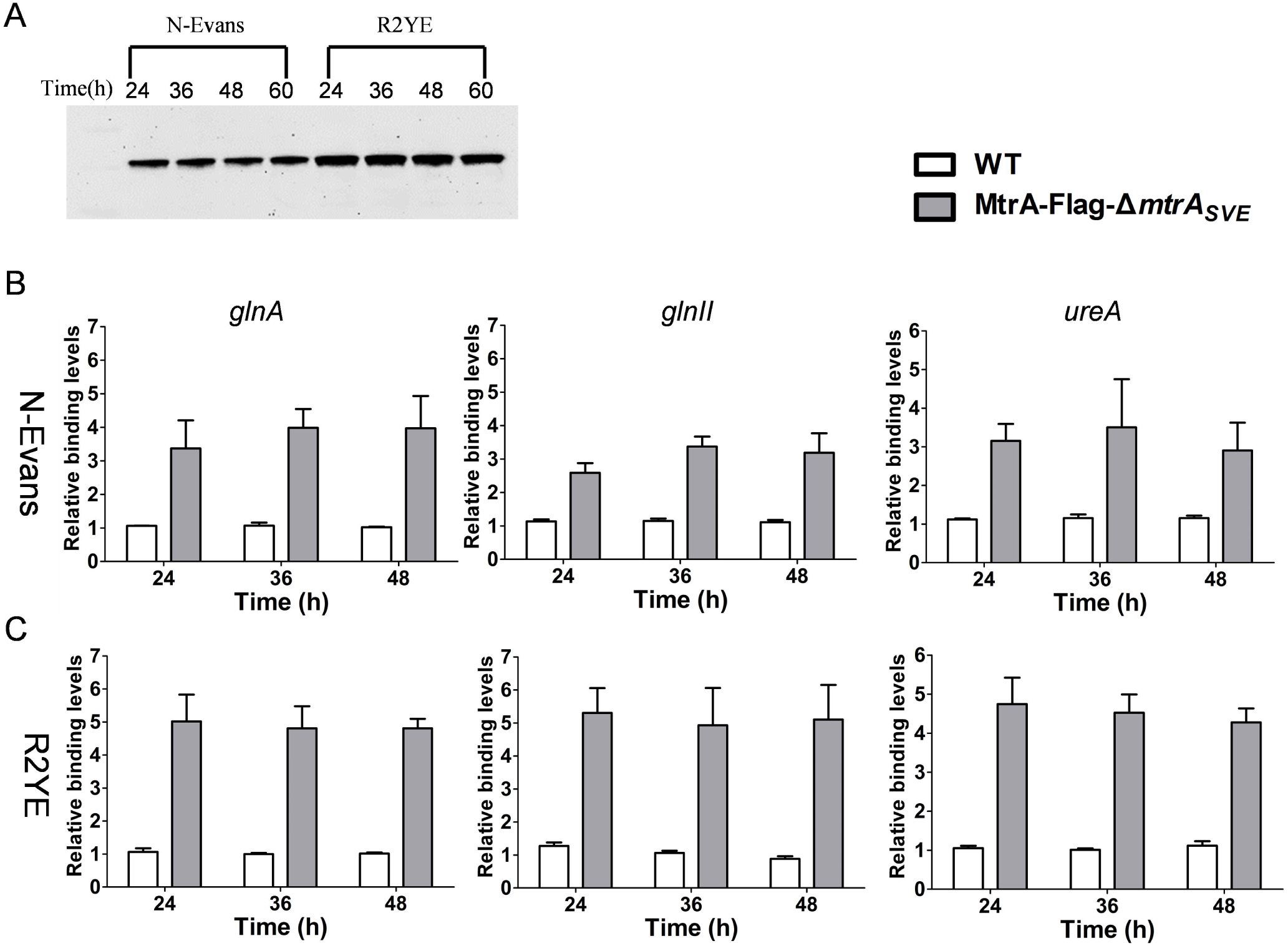
Comparison of the level of MtrA and its binding to the promoters of nitrogen metabolism genes under nitrogen-limited and nutrient-rich conditions. (A) Western blot analysis using 10 μg total cellular lysates extracted at indicated times from mycelia of indicated strains grown on R2YE or N-Evans. (B-C) ChIP-qPCR analysis of the binding of MtrA to the promoters of nitrogen metabolism genes from cultures grown on R2YE or N-Evans. Analysis was performed using strains MtrA-FLAG-Δ*mtrA*_*SVE*_ and the wild-type 10712 (WT) grown on R2YE or N-Evans for the indicated times. The y-axis shows the binding levels of MtrA-FLAG in MtrA-FLAG-Δ*mtrA*_*SVE*_ and WT relative to background levels, which was determined by recovery of target sequences. Results are the means (±SD) of triplet biological experiments.

### GlnR binds differentially to the MtrA/GlnR sites of nitrogen metabolism genes under different growth conditions

To investigate if the differential expression of *glnR* on different media is reflected at the protein level, we performed Western blot analysis using the complemented strain GlnR-FLAG-Δ*glnR*_*SVE*_, which expresses a functional GlnR-FLAG (Fig. S12). Crude cellular lysates were extracted at the same four time points of GlnR-FLAG-Δ*glnR*_*SVE*_ grown on solid R2YE and N-Evans. Although the level of GlnR was nearly constant on either R2YE or N-Evans, the level on N-Evans was notably higher than on R2YE (Fig. 7A), consistent with the transcriptional analysis (Fig. 5). To determine whether the different levels of GlnR leads to differential binding *in vivo* to the MtrA/GlnR sites for nitrogen metabolism genes, we performed ChIP-qPCR analysis and compared the binding level of GlnR to nitrogen genes on R2YE and N-Evans (Fig. 7B, 7C and Fig. S20). Higher binding levels were detected on N-Evans than on R2YE and the levels are fairly constant for a given medium. For example, at 24 h, the levels on N-Evans versus on R2YE were as follows: *glnA* (5.71±0.69) vs 4.35±0.69), *glnII* (4.77±0.60 vs 3.63±0.64), *ureA* (4.15±0.71 vs 3.24±0.52), *amtB* (4.76±0.35 vs 3.22±0.19)], and several other nitrogen genes (Fig. S20), suggesting that GlnR binds stronger under nitrogen-limited conditions than under nutrient-rich conditions.

**Figure 7.**
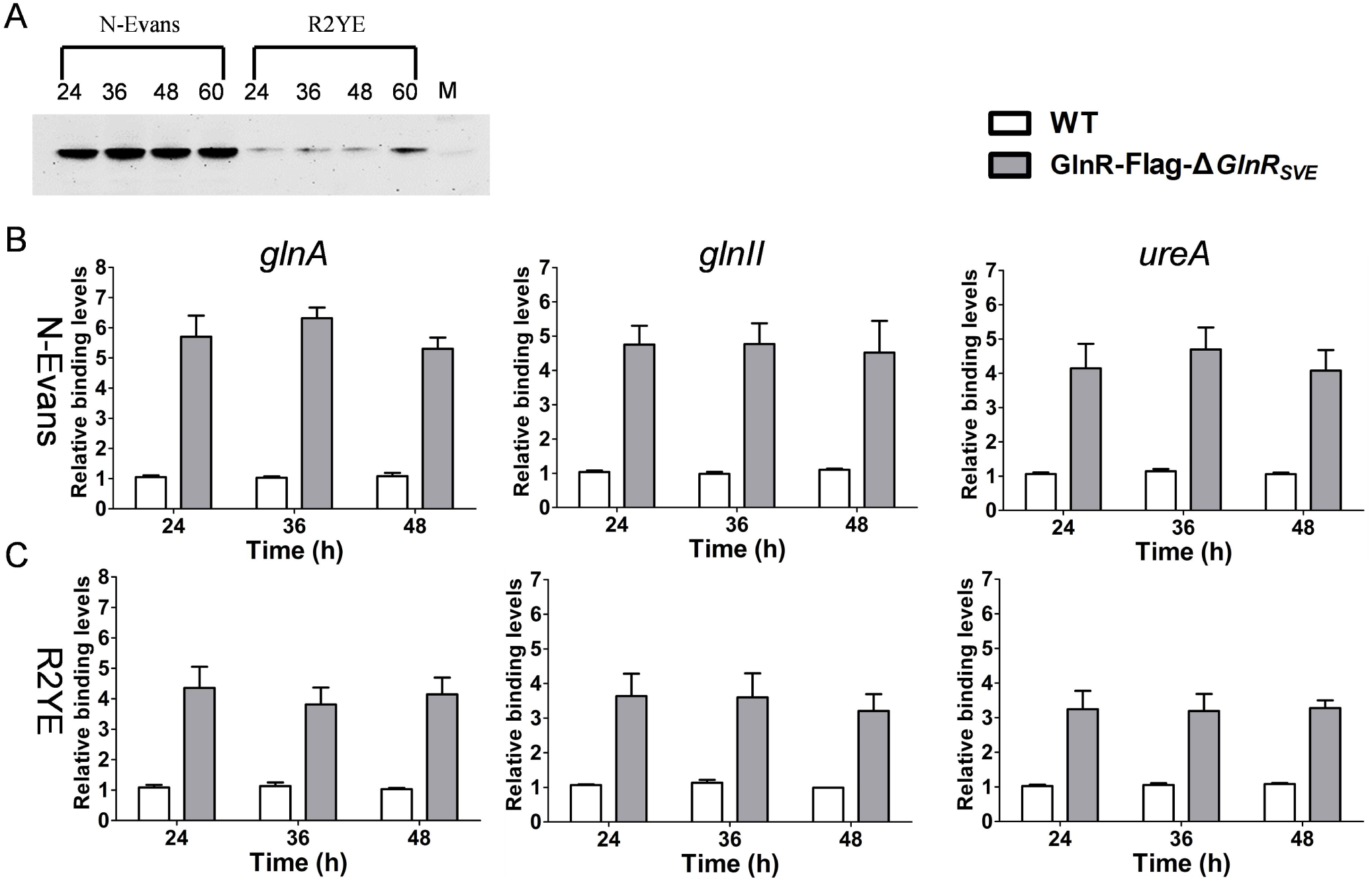
Comparison of the level of GlnR and its binding to the promoters of nitrogen metabolism genes under nitrogen-limited and nutrient-rich conditions. (A) Western blot analysis. For details, see legend to Figure 6, panel A. (B-C) ChIP-qPCR analysis of the binding of GlnR to the promoters of nitrogen metabolism genes from cultures grown on R2YE or N-Evans. Analysis was performed using strains GlnR-FLAG-Δ*glnR*_*SVE*_, and the wild-type 10712 (WT) grown on R2YE for N-Evans for the indicated times. The y-axis shows the binding levels of GlnR-FLAG in GlnR-FLAG-Δ*glnR*_*SVE*_ and WT relative background levels, which was determined by recovery of target sequences. Results are the means (±SD) of triplet biological experiments.

## Discussion

From the model that we proposed for the regulation of nitrogen metabolism genes (Zhu *et al*., 2019), MtrA binds the MtrA sites of nitrogen assimilation genes and represses these genes to prevent the unnecessary expression of these genes when nitrogen resources are in surplus under nutrient-rich conditions; however, GlnR binds the GlnR boxes of these genes and activates the genes under nitrogen-limited conditions. As there is strong similarity in the core sequence of the MtrA site and the GlnR box and as MtrA and GlnR compete *in vitro* to bind to these sequences (Zhu *et al*., 2019), we hypothesized that MtrA would bind the MtrA sites/GlnR boxes under nutrient-rich conditions and disassociate from them under nitrogen-limited conditions *in vivo*, whereas GlnR would exhibit the opposite pattern, with binding under nitrogen-limited conditions and disassociation under nutrient-rich conditions. However, the hypothesis was partially contradicted by the data obtained from this study. We showed that MtrA and GlnR bound the MtrA sites/GlnR boxes under both nitrogen-limited and nutrient-rich conditions, although the binding levels differed. In general, MtrA bound more strongly to these target sites under nutrient-rich conditions than under nitrogen-limited conditions, in agreement with a moderately higher levels of MtrA under nutrient-rich conditions than under nitrogen-limited conditions. However, GlnR bound at notably stronger levels to the MtrA sites/GlnR boxes under nitrogen-limited conditions than under nutrient-rich conditions, consistent with a markedly higher level of GlnR under nitrogen-limited conditions. Although it has only a minor role in the regulation of nitrogen metabolism genes under nitrogen-limited conditions, MtrA still bound the MtrA sites/GlnR boxes under these conditions; likewise, GlnR still bound these targets under nutrient-rich conditions, implying co-occupancy of the MtrA sites/GlnR boxes by MtrA and GlnR under the conditions tested. However, when both proteins are bound to the same site, it is unclear how MtrA exerts its role under nutrient-rich conditions and GlnR exerts its role under nitrogen-limited conditions. Nevertheless, as MtrA demonstrated a higher binding level than GlnR under nutrient-rich conditions, MtrA may occupy more of the MtrA sites/GlnR boxes than GlnR does under nutrient-rich conditions, enabling MtrA to manifest its repressor role when nitrogen sources are abundant. In contrast, as GlnR generally displayed a higher binding level than MtrA did under nitrogen-limited conditions, GlnR may occupy more of these sites when nitrogen is limited, consistent with its role as an activator of nitrogen metabolism genes.

The lower level of MtrA under nitrogen-limited and of GlnR under nutrient-rich conditions conditions could be caused at the transcriptional level and potentially at the post-transcriptional level. In addition to changes in the levels of these regulators, modification at the post-translational level has been reported for GlnR in *Streptomyces* and MtrA in *Mycobacterium tuberculosis* (Singh *et al*., 2020, Singhal *et al*., 2020, Amin *et al*., 2016). Acetylated and phosphorylated forms of GlnR have been identified (Amin *et al*., 2016); GlnR phosphorylation correlated with nitrogen-rich conditions, and phosphorylation inhibited the binding of GlnR to its target genes, whereas acetylation had only a minor influence on the binding of GlnR to its target genes (Amin *et al*., 2016). Acetylation and methylation of MtrA influenced its repressor activity in *M. tuberculosis* (Singh *et al*., 2020, Singhal *et al*., 2020), and MtrA of *Streptomyces* may be similarly subject to post-translational modification, with different forms of MtrA having different binding affinities for target genes (Singh *et al*., 2020, Singhal *et al*., 2020). However, the role of any such modifications needs to be further investigated in *Streptomyces*.

The transcriptional data obtained from mutant strains with deletion of a single gene (Δ*mtrA* or Δ*glnR*) in this and a previous study (Zhu *et al*., 2019) indicated that MtrA and GlnR function under nutrient-rich and nitrogen-limited conditions, respectively. Therefore, we initially hypothesized that, under nutrient-rich conditions, the transcriptional pattern of nitrogen metabolism genes in the double mutant Δ*mtrA*-*glnR* would follow the pattern of Δ*mtrA*, while under nitrogen-limited conditions, the pattern would follow that of Δ*glnR*. Consistent with our hypothesis, the transcriptional pattern in Δ*mtrA*-*glnR* was similar to that of Δ*glnR* under nitrogen-limited conditions (Fig. 1B). However, unexpectedly, the transcriptional pattern in Δ*mtrA*-*glnR* was also similar to that of Δ*glnR* under nutrient-rich conditions (Fig. 1A), implying a GlnR-dependent regulatory effect of MtrA, although GlnR maintains a lower level of expression under this condition. This is a new and interesting finding, although it is difficult to explain from our current understanding.

Based on our data, we propose a new model for nitrogen regulation by MtrA and GlnR (Fig. 8). In this model, both MtrA and GlnR are auto-regulatory, repressing their own expression. Under nitrogen-poor conditions, expression of GlnR is highly induced, and therefore GlnR binds more strongly (or more GlnR binds) to the MtrA sites/GlnR boxes of the nitrogen metabolism genes, and thus these genes are activated. Under nutrient-rich conditions, expression of GlnR is minimal whereas MtrA is induced, and therefore MtrA binds more strongly (or more MtrA binds) to the target sites, resulting in repression of the nitrogen genes. The molecular mechanism underlying the dependence of MtrA on GlnR for the regulation of nitrogen metabolism genes is not clear and is not yet explained in our model. Nevertheless, our study does provide new insights into the understanding of the complex regulation of nitrogen metabolism in microbes.

**Figure 8.**
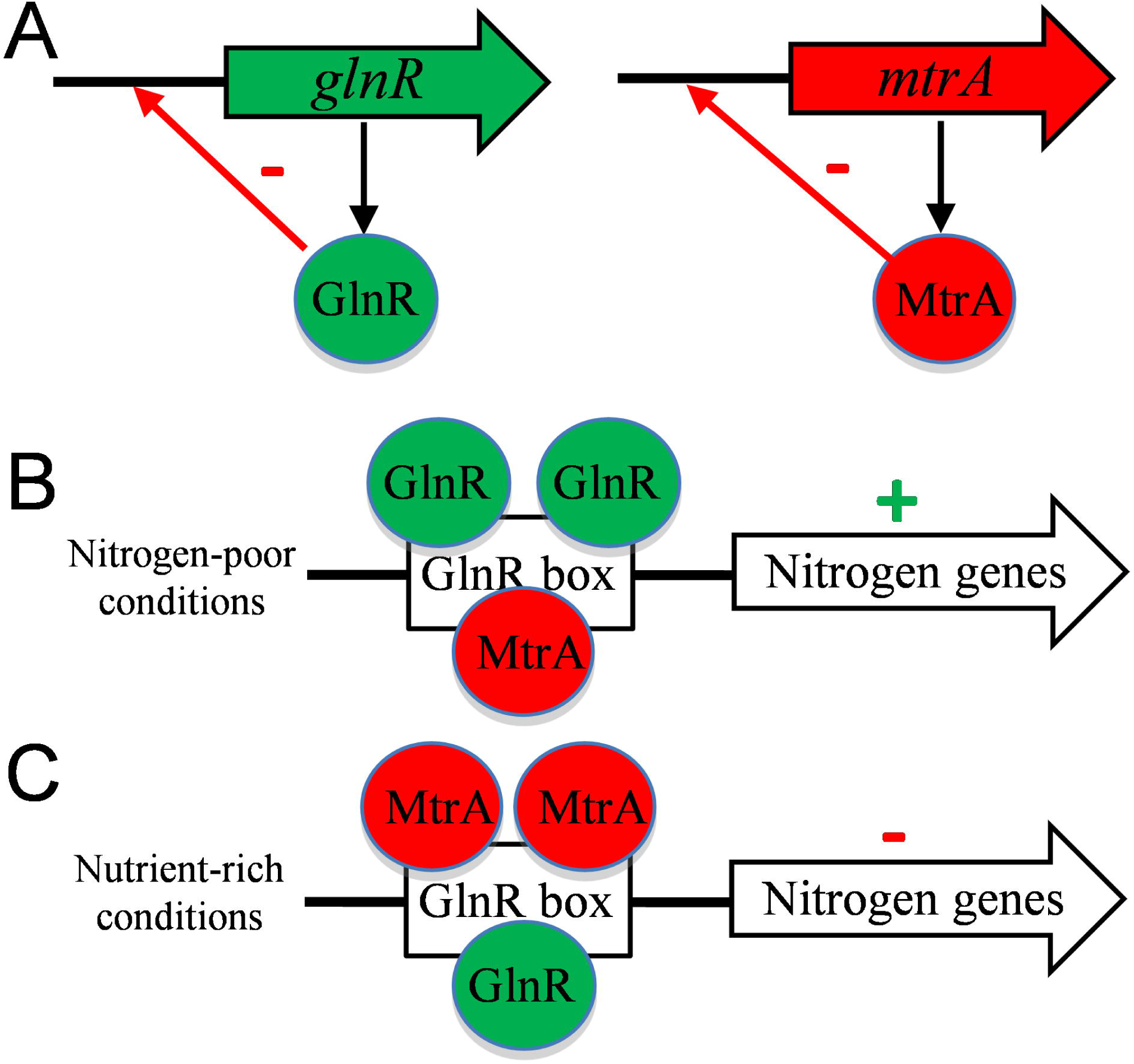
Model for regulation of nitrogen metabolism genes by MtrA and GlnR. (A) MtrA and GlnR are auto-regulatory, repressing their own expression. (B) Under nitrogen-poor conditions, more GlnR binds to the MtrA sites/GlnR boxes, activating the nitrogen metabolism genes. (C) Under nutrient-rich conditions, more MtrA binds to the MtrA sites/GlnR boxes, repressing the nitrogen metabolism genes.

### Experimental procedures

#### Strains, plasmids, primers, and culture conditions

All strains and primers are listed in Table S1 and Table S2, respectively. *Streptomyces venezuelae* ATCC 10712 was used as the wild-type strain in this study. The *Streptomyces* strains were cultivated on solid maltose-yeast extract-malt extract (MYM) medium (Frojd & Flardh, 2019) for sporulation, and on N-Evans (Fink *et al*., 2002), YBP (Ou *et al*., 2009), and R2YE (Kieser *et al*., 2000) for phenotypic observation, RNA extraction, cellular lysate purification, and ChIP analysis. All *Escherichia coli* strains were cultured in Luria-Bertani (LB) agar or liquid medium. When necessary, appropriate antibiotics were added.

#### Deletion of both mtrA and glnR from the genome of S. venezuelae

The mutant strain Δ*mtrA*-*glnR* with deletions of both *mtrA and glnR* was obtained using the mutation plasmid pJTU-*mtrA* to delete *mtrA* from the mutant strain Δ*glnR* (Zhu *et al*., 2020a). Plasmid pJTU-*mtrA*, which is apramycin resistance (Zhu *et al*., 2020a), was transformed into *E. coli* ET12567 (pUZ8002) and then introduced into the kanamycin-resistant Δ*glnR*_*SVE*_ (Zhu *et al*., 2020a) by conjugation. After several rounds of selection on MS agar containing both apramycin and kanamycin, the deletion of *mtrA* from Δ*glnR*_*SVE*_ was confirmed by PCR using MtrA V-F/R.

#### Expression and purification of MtrA and GlnR

His-tagged MtrA and GlnR were expressed and purified essentially as described (Zhu *et al*., 2020a, Zhu *et al*., 2019, Lu *et al*., 2018). In brief, protein production was induced by addition of 1 mM IPTG, and bacterial cells were collected after overnight culture at 16°C and then re-suspended and sonicated in binding buffer [50 mM NaH_2_PO_4_ (pH 8.0), 200 mM NaCl, 20 mM imidazole] on ice. Crude lysates were centrifuged to remove cell debris, and soluble proteins in supernatant were purified by Ni affinity column (Qiagen, USA). Purified proteins were examined by sodium dodecyl sulfate-polyacrylamide gel electrophoresis (SDS-PAGE), and their concentration was determined by the Pierce BCA Protein Assay Kit (Thermo Scientific, USA).

#### Electrophoretic mobility shift assays (EMSAs)

All primers for EMSAs (Table S2) were labelled with biotin at the 5’-terminus. The complementary forward and reverse 59 nt primers were mixed and annealed to produce probes. The conditions for EMSAs were as described previously (Zhang *et al*., 2015, Zhu *et al*., 2020b). Signal detection was conducted by the ECL Western Blotting Analysis System kit (GE Healthcare) and was displayed by exposure to X-ray film or visualized by myECL imager (Thermo Scientific) instrument.

#### Extraction of crude cellular lysates, SDS-PAGE, and Western blot analysis

*S. venezuelae* strains were cultivated on solid R2YE or N-Evans (supplemented with 2 mM glutamine), and mycelia were harvested at indicated times. The harvested mycelia were ground in liquid nitrogen, dissolved in lysis buffer (50 mM Tris-HCl, 50 mM EDTA, PH8.0), and centrifuged to remove cellular debris. The concentration of the crude lysates was determined by the Pierce Protein Assay Kit. Equal amounts of crude lysates were separated by SDS-PAGE (12%) and then transferred to Hybond-ECL membranes (GE Amersham), which were blocked with 5% fat-free milk at room temperature for 2 hours, washed twice, and incubated with anti-FLAG mAb (1:3000; Boster Biological Technology) at 4°C overnight (Yan *et al*., 2020, Lu *et al*., 2020b). Next, the membranes were washed twice before incubating with the HRP-conjugated goat anti-mouse IgG (H+L) secondary antibody (1:5000; Boster Biological Technology) for 50 min at room temperature. Finally, the membranes were washed twice, and the signal was revealed by the ECL Western Blotting Analysis System kit or imaged by the myECL imager system.

#### Total RNA extraction, reverse transcription-PCR (RT-PCR), and real-time PCR

Equal amounts of spores of *Streptomyces* strains were inoculated onto solid YBP, R2YE, and N-Evans media, and cultures were collected at indicated times. For the media-shift experiment, the *Streptomyces* strains were first cultured in liquid YBP medium for 24 h at 30.0 °C and at 220 rpm. After the optical density of the culture reached 2.0 at OD_450nm_, one portion of the culture was centrifuged and collected for RNA isolation at the base time (0 h). An equal portion of the YBP culture was centrifuged, washed twice with liquid N-Evans, and dispersed into 50 ml N-Evans medium supplemented with 2 mM glutamine for extended growth (4 h or 6 h) at 30.0°C and at 220 rpm. Cell cultures were collected, ground in liquid nitrogen, and processed for RNA extraction as described previously (Zhang *et al*., 2017, Lu *et al*., 2020a). Reverse transcription PCR for cDNA synthesis and real-time PCR assays were carried out as described previously (Zhu *et al*., 2020c). Specificity and melting curves of the PCR products were determined using the Roche LightCycler480 thermal cycler according to the manufacturer’s protocol. Transcription levels of measured genes were normalized relative to the level for *hrdB*, which was used as the internal control.

#### Construction of engineered strains expressing FLAG-tagged MtrA or GlnR

To express the MtrA-FLAG fusion protein, the plasmid pMtrA-FLAG was constructed following the described strategy (Liu *et al*., 2019). Briefly, DNA fragment I containing the promoter and coding region of *mtrA* of *S. venezuelae* was amplified using the primer set MtrA Fcom-F/R and the template genomic DNA, and DNA fragment II containing the linker and 3 × FLAG sequence (including stop codon) was amplified using primer set Linker-Flag-F/R with the template pMacR-FLAG, which contains the linker sequence and the coding sequence of FLAG (Liu *et al*., 2019); the two sets of primers were designed so that there would be overlapping sequences between these two amplified fragments. The two PCR fragments were purified, mixed, and ligated with pMS82 to obtain pMtrA-FLAG, which was then introduced into Δ*mtrA*_*SVE*_ and Δ*mtrA-glnR* by conjugation to obtain the complemented strains MtrA-Flag-Δ*mtrA*_*SVE*_ and MtrA-Flag-Δ*mtrA-glnR*, respectively. The plasmid expressing the GlnR-FLAG fusion protein and complemented strains GlnR-Flag-Δ*glnR*_*SVE*_ and GlnR-Flag-Δ*mtrA-glnR* were constructed similarly, using primers listed in Table S2.

#### Chromatin immunoprecipitation and qPCR

*S. venezuelae* strains were grown on R2YE and N-Evans agar and harvested at indicated times. For the chromatin immunoprecipitation (ChIP), the M2 mouse monoclonal anti-FLAG antibody (Sigma) was used. The cross-linking, chromosomal DNA sonication, immunoprecipitation, reverse of the cross-links, and elution steps were conducted essentially as described previously (Bush *et al*., 2013, Liu *et al*., 2019, Bush *et al*., 2019). The elution was quantified and subjected to qPCR analysis. The qPCR reactions were performed as above. To calculate the binding level of protein in the ChIP samples, the relative quantities of each DNA fragment were normalized with the housekeeping gene *hrdB*, which served as an internal control, and the binding level of at each target in the input chromosomal DNA was arbitrarily set to one.

## Acknowledgement

This work was supported by grants from the National Natural Science Foundation of Shandong Province (ZR2019MC062 to XP), the Open Funding Project of the State Key Laboratory of Microbial Metabolism (MMLKF21-02 to XP), and the National Key R&D Program of China (2018YFA0900400 to AL).

## Conflict of interest

The authors declare that they have no conflict of interest with the contents of this article.

## Author contributions

XP conceived, supervised the study, and wrote the paper; XP and YZ designed experiments; YZ, JW, TL, and WS performed experiments; XP, YZ, and AL analysed data; and all authors reviewed the results and approved the final version of the manuscript.

## Notes

### Competing Interest Statement

The authors have declared no competing interest.

## References

Amin, R., Franz-Wachtel, M., Tiffert, Y., Heberer, M., Meky, M., Ahmed, Y., et al. (2016) Post-translational Serine/Threonine Phosphorylation and Lysine Acetylation: A Novel Regulatory Aspect of the Global Nitrogen Response Regulator GlnR in S. coelicolor M145. Frontiers in molecular biosciences 3: 38.

Bush, M.J., Bibb, M.J., Chandra, G., Findlay, K.C., and Buttner, M.J. (2013) Genes required for aerial growth, cell division, and chromosome segregation are targets of WhiA before sporulation in Streptomyces venezuelae. Mbio 4: e00684–00613.

Bush, M.J., Chandra, G., Al-Bassam, M.M., Findlay, K.C., and Buttner, M.J. (2019) BldC Delays Entry into Development To Produce a Sustained Period of Vegetative Growth in Streptomyces venezuelae. MBio 10.

Chater, K., (2011) Differentiation in Streptomyces: the properties and programming of diverse cell-types. In: Streptomyces: Molecular Biology and Biotechnology. D. P (ed). Caister Academic Press, pp. 43–86.

Fink, D., Weissschuh, N., Reuther, J., Wohlleben, W., and Engels, A. (2002) Two transcriptional regulators GlnR and GlnRII are involved in regulation of nitrogen metabolism in Streptomyces coelicolor A3(2). Mol Microbiol 46: 331–347.

Frojd, M.J., and Flardh, K. (2019) Apical assemblies of intermediate filament-like protein FilP are highly dynamic and affect polar growth determinant DivIVA in Streptomyces venezuelae. Mol Microbiol 112: 47–61.

Hopwood, D.A., (2007) Streptomyces in Nature and Medicine. In.: OXFORD UNIVERSITY PRESS, pp.

Kawamoto, S., and Ochi, K. (1998) Comparative ribosomal protein (L11 and L30) sequence analyses of several Streptomyces spp. commonly used in genetic studies. Int J Syst Bacteriol 48 Pt 2: 597–600.

Kieser, T., Bibb, M.J., Buttner, M.J., Chater, K.F., and Hopwood, D.A., (2000) Practical Streptomyces Genetics. In.: Norwich: John Innes Foundation., pp.

Leigh, J.A., and Dodsworth, J.A. (2007) Nitrogen regulation in bacteria and archaea. Annu Rev Microbiol 61: 349–377.

Lewis, R.A., Laing, E., Allenby, N., Bucca, G., Brenner, V., Harrison, M., et al. (2010) Metabolic and evolutionary insights into the closely-related species Streptomyces coelicolor and Streptomyces lividans deduced from high-resolution comparative genomic hybridization. BMC Genomics 11: 682.

Liu, M., Zhang, P., Zhu, Y., Lu, T., Wang, Y., Cao, G., et al. (2019) Novel Two-Component System MacRS Is a Pleiotropic Regulator That Controls Multiple Morphogenic Membrane Protein Genes in Streptomyces coelicolor. Appl Environ Microbiol 85.

Lu, T., Cao, Q., Pang, X.H., Xia, Y.Z., Xun, L.Y., and Liu, H.W. (2020a) Sulfane sulfur-activated actinorhodin production and sporulation is maintained by a natural gene circuit in Streptomyces coelicolor. Microb Biotechnol 13: 1917–1932.

Lu, T., Zhu, Y.P., Zhang, P.P., Sheng, D.H., Cao, G.X., and Pang, X.H. (2018) SCO5351 is a pleiotropic factor that impacts secondary metabolism and morphological development in Streptomyces coelicolor. Fems Microbiol Lett 365.

Lu, X.R., Liu, X.C., Chen, Z., Li, J.L., van Wezel, G.P., Chen, W., et al. (2020b) The ROK-family regulator Rok7B7 directly controls carbon catabolite repression, antibiotic biosynthesis, and morphological development in Streptomyces avermitilis. Environ Microbiol 22: 5090–5108.

Merrick, M.J., and Edwards, R.A. (1995) Nitrogen control in bacteria. Microbiological reviews 59: 604–622.

Ou, X.J., Zhang, B., Zhang, L., Zhao, G.P., and Ding, X.M. (2009) Characterization of rrdA, a TetR Family Protein Gene Involved in the Regulation of Secondary Metabolism in Streptomyces coelicolor. Appl Environ Microb 75: 2158–2165.

Pullan, S.T., Chandra, G., Bibb, M.J., and Merrick, M. (2011) Genome-wide analysis of the role of GlnR in Streptomyces venezuelae provides new insights into global nitrogen regulation in actinomycetes. BMC Genomics 12: 175.

Rodriguez-Garcia, A., Sola-Landa, A., Apel, K., Santos-Beneit, F., and Martin, J.F. (2009) Phosphate control over nitrogen metabolism in Streptomyces coelicolor: direct and indirect negative control of glnR, glnA, glnII and amtB expression by the response regulator PhoP. Nucleic Acids Res 37: 3230–3242.

Singh, K.K., Athira, P.J., Bhardwaj, N., Singh, D.P., Watson, U., and Saini, D.K. (2020) Acetylation of Response Regulator Protein MtrA in M. tuberculosis Regulates Its Repressor Activity. Frontiers in microbiology 11: 516315.

Singhal, A., Virmani, R., Naz, S., Arora, G., Gaur, M., Kundu, P., et al. (2020) Methylation of two-component response regulator MtrA in mycobacteria negatively modulates its DNA binding and transcriptional activation. Biochem J 477: 4473–4489.

Som, N.F., Heine, D., Holmes, N., Knowles, F., Chandra, G., Seipke, R.F., et al. (2017a) The MtrAB two-component system controls antibiotic production in Streptomyces coelicolor A3(2). Microbiology 163: 1415–1419.

Som, N.F., Heine, D., Holmes, N.A., Munnoch, J.T., Chandra, G., Seipke, R.F., et al. (2017b) The Conserved Actinobacterial Two-Component System MtrAB Coordinates Chloramphenicol Production with Sporulation in Streptomyces venezuelae NRRL B-65442. Frontiers in microbiology 8: 1145.

Tiffert, Y., Franz-Wachtel, M., Fladerer, C., Nordheim, A., Reuther, J., Wohlleben, W., et al. (2011) Proteomic analysis of the GlnR-mediated response to nitrogen limitation in Streptomyces coelicolor M145. Appl Microbiol Biotechnol 89: 1149–1159.

Tiffert, Y., Supra, P., Wurm, R., Wohlleben, W., Wagner, R., and Reuther, J. (2008) The Streptomyces coelicolor GlnR regulon: identification of new GlnR targets and evidence for a central role of GlnR in nitrogen metabolism in actinomycetes. Mol Microbiol 67: 861–880.

Wang, J., Wang, Y., and Zhao, G.P. (2015) Precise characterization of GlnR Box in actinomycetes. Biochem Biophys Res Commun 458: 605–607.

Wang, R., Mast, Y., Wang, J., Zhang, W., Zhao, G., Wohlleben, W., et al. (2013) Identification of two-component system AfsQ1/Q2 regulon and its cross-regulation with GlnR in Streptomyces coelicolor. Mol Microbiol 87: 30–48.

Wolfgang Wohlleben, Y.M.a.J.R., (2011) Regulation of nitrogen assimilation in Streptomycetes and other actinobacteria. In: Streptomyces-Molecular Biology and Biotechnologh. P. Dyson (ed). Caister Academic Press, pp.

Yan, H., Lu, X.R., Sun, D., Zhuang, S., Chen, Q., Chen, Z., et al. (2020) BldD, a master developmental repressor, activates antibiotic production in two Streptomyces species. Mol Microbiol 113: 123–142.

Zhang, P., Wu, L., Zhu, Y., Liu, M., Wang, Y., Cao, G., et al. (2017) Deletion of MtrA Inhibits Cellular Development of Streptomyces coelicolor and Alters Expression of Developmental Regulatory Genes. Frontiers in microbiology 8: 2013.

Zhang, P.P., Zhao, Z.L., Li, H., Chen, X.L., Deng, Z.X., Bai, L.Q., et al. (2015) Production of the antibiotic FR-008/candicidin in Streptomyces sp FR-008 is co-regulated by two regulators, FscRI and FscRIV, from different transcription factor families. Microbiol-Sgm 161: 539–552.

Zhu, Y., Zhang, P., Lu, T., Wang, X., Li, A., Lu, Y., et al. (2021) Impact of MtrA on phosphate metabolism genes and the response to altered phosphate conditions in Streptomyces. Environ Microbiol.

Zhu, Y., Zhang, P., Zhang, J., Wang, J., Lu, Y., and Pang, X. (2020a) Impact on Multiple Antibiotic Pathways Reveals MtrA as a Master Regulator of Antibiotic Production in Streptomyces spp. and Potentially in Other Actinobacteria. Appl Environ Microbiol 86.

Zhu, Y., Zhang, P., Zhang, J., Xu, W., Wang, X., Wu, L., et al. (2019) The developmental regulator MtrA binds GlnR boxes and represses nitrogen metabolism genes in Streptomyces coelicolor. Mol Microbiol 112: 29–46.

Zhu, Y.P., Lu, T., Zhang, J., Zhang, P.P., Tao, M.F., and Pang, X.H. (2020b) A novel XRE family regulator that controls antibiotic production and development in Streptomyces coelicolor. Appl Microbiol Biot 104: 10075–10089.

Zhu, Y.P., Xu, W.H., Zhang, J., Zhang, P.P., Zhao, Z.L., Sheng, D.H., et al. (2020c) A hierarchical network of four regulatory genes controlling production of the polyene antibiotic candicidin in Streptomyces sp. Strain FR-008. Appl Environ Microb 86: e00055–00020.

