## supplemental TableS1-S2 FigureS1-S20 for "Effects of dual deletion of *glnR* and *mtrA* on expression of nitrogen metabolism genes in *Streptomyces venezuelae*"

Table S1. Bacterial strains and plasmids used in this study

| Strain or plasmid | Description | Reference |
|---|---|---|
| **Strains** | | |
| ***Streptomyces*** | | |
| *S. venezuelae* ATCC 10712 | Wild type | (He, et al., 2016) |
| Δ*mtrA*$_{SVE}$ | *mtrA* deletion strain, Apr$^R$ | (Zhu, et al., 2020) |
| C-Δ*mtrA*$_{SVE}$ | Δ*mtrA* complemented with pCom-mtrA, Apr$^R$, Hyg$^R$ | This study |
| MtrA-Flag-Δ*mtrA*$_{SVE}$ | Δ*mtrA* complemented with pMtrA-Flag, Apr$^R$, Hyg$^R$ | This study |
| Δ*glnR*$_{SVE}$ | *glnR* deletion strain, Km$^R$ | (Zhu, et al., 2020) |
| C-Δ*glnR*$_{SVE}$ | Δ*glnR* complemented with pCom-glnR, Km$^R$, Hyg$^R$ | This study |
| GlnR-Flag-Δ*glnR*$_{SVE}$ | Δ*glnR* complemented with pGlnR-Flag, Km$^R$, Hyg$^R$ | This study |
| Δ*mtrA-glnR* | *mtrA* and *glnR* deletion strain, Apr$^R$, Km$^R$ | This study |
| ***E. coli*** | | |
| DH5α | Strain use for general cloning | Shanghai Weidi |
| BL21 (DE3) | Strain used for protein expression, Cm$^R$ | Sangon |
| ET12567(pUZ8002) | Strain used for conjugation between *E.coli* and *Streptomyces*, containing helper plasmid pUZ8002, Cm$^R$, Km$^R$ | (Kieser, et al., 2000) |
| **Plasmids** | | |
| pMD18-T | General cloning vector, Amp$^R$ | Takara |
| pJTU1278 | Shuttle vector for conjugation,Amp$^R$ | (He, et al., 2010) |
| pMS82 | *Streptomyces* integrative vector, Hyg$^R$ | (Gregory, et al., 2003) |
| pIJ773 | Plasmid used for cloning aparmycin resistance genes, Apr$^R$ | Norwich, UK |
| pET15b | Plasmid used for protein expression in *E. coli*, Amp$^R$ | Novagen |
| pET28a | Plasmid used for protein expression in *E. coli*, Km$^R$ | Novagen |
| pJTU-mtrA | pJTU1278 with left (2108-bp), aparmycin resistance genes and right (2071-bp) flanking sequences of *mtrA*, Amp$^R$, Apr$^R$ | This study |
| pJTU-glnR | pJTU1278 with left (1383-bp), kanamycin resistance genes and right (1349-bp) flanking sequences of *glnR*, Amp$^R$, Km$^R$ | This study |
| pCom-mtrA | pMS82 with 880-bp upstream sequence and 678-bp coding sequence of *mtrA*, Hyg$^R$ | This study |
| pCom-glnR | pMS82 with 382-bp upstream sequence and 831-bp coding sequence of *glnR*, Hyg$^R$ | This study |
| pMtrA-Flag | a 3 ×FLAG epitope inserted before the stop codon of *mtrA*, Hyg$^R$ | This study |
| pGlnR-Flag | a 3 ×FLAG epitope inserted before the stop codon of *glnR*, Hyg$^R$ | This study |
| pEASY-BLUNT | General cloning vector, Amp$^R$, Km$^R$ | Takara |
| pEX-*glnR*$_{SVE}$ | *glnR* (of *S. venezuelae*) expression plasmid, Km$^R$, Cm$^R$ | (Zhu, et al., 2020) |
| pEX-*mtrA*$_{SVE}$ | *mtrA* (of *S. venezuelae*) expression plasmid, Amp$^R$, Cm$^R$ | (Zhu, et al., 2020) |

### Reference:


Gregory, M.A., Till, R., and Smith, M.C.M. (2003) Integration site for streptomyces phage phi BT1 and development of site-specific integrating vectors, *J Bacteriol* **185**: 5320-5323.

He, J.M., Zhu, H., Zheng, G.S., Liu, P.P., Wang, J., Zhao, G.P., et al. (2016) Direct involvement of the master nitrogen metabolism regulator GlnR in antibiotic biosynthesis in Streptomyces, *J Biol Chem* **291**: 26443-26454.

He, Y.L., Wang, Z.J., Bai, L.Q., Liang, J.D., Zhou, X.F., and Deng, Z.X. (2010) Two pHZ1358 derivative vectors for

efficient gene knockout in Streptomyces, *J Microbiol Biotechn* **20**: 678-682.
Kieser, T., Bibb, M.J., Buttner, M.J., Chater, K.F., and Hopwood, D.A. (eds) (2000) *Practical Streptomyces Genetics*: Norwich: John Innes Foundation
Zhu, Y.P., Zhang, P.P., Zhang, J., Wang, J., Lu, Y.H., and Pang, X.H. (2020) Impact on multiple antibiotic pathways reveals MtrA as a master regulator of antibiotic production in Streptomyces spp. and potentially in other Actinobacteria, *Appl Environ Microb* **86**: e01201-01220.

Table 2. Primers used in this study

| Primers | Sequence (5′-3′) |
|---|---|
| **For construction of strain expressing MtrA-FLAG** | |
| MtrA comF | AAGCTTGTCGAGGGACTGCTCCAGGGCCTC (HindIII) |
| MtrA comR | AAGCTTGGGCTTCGGAGCAGCACTGCCTGT (HindIII) |
| MtrA Fcom-F | ACCCGGGGATCCTCTAGAGATTGTCGAGGGACTGCTCCAGGGCCTC |
| MtrA Fcom-R | CCGCCTGAACCGCCTCCACCGCTCGGTCCCGCCTTGTACCCGACACCA |
| LinkerFlag-F | GGTGGAGGCGGTTCAGGCGGAGG |
| LinkerFlag-R | GCATGCCTGCAGGTCGACGATATCACTTGTCATCGTCATCCT |
| **For construction of strain expressing GlnR-FLAG** | |
| GlnR comF | AAGCTTCGGAGCAGGGTCGTAAACCGCCTCG (HindIII) |
| GlnR comR | AAGCTTGCCCAGTTCCTTGATCGCGGCGAGG (HindIII) |
| GlnR Fcom-F | ACCCGGGGATCCTCTAGAGATTCGGAGCAGGGTCGTAAACCGCCTCG |
| GlnR Fcom-R | CCGCCTGAACCGCCTCCACCCCTACCGGCAGGTCGCACTGTGGCGT |
| **For EMSA analysis** | |
| SVEN0835 WT F | CATGAGATCGCAAAGCCGCAGTTAACGAGCCGGAAAAACTTCCGCCCTAACGTGCAATG |
| SVEN0835 WT R | CATTGCACGTTAGGGCGGAAGTTTTTCCGGCTCGTTAACTGCGGCTTTGCGATCTCATG |
| SVEN0835 M1 F | CATGAGATCGCAAAGCCGCAGTCAGCGAGCCGAGGGAACTTCCGCCCTAACGTGCAATG |
| SVEN0835 M1 R | CATTGCACGTTAGGGCGGAAGTTCCCTCGGCTCGCTGACTGCGGCTTTGCGATCTCATG |
| SVEN0835 M2 F | CATGAGATCGCAAAGCCGCAGTTAACGCCTCTGAAAAACTTCCGCCCTAACGTGCAATG |
| SVEN0835 M2 R | CATTGCACGTTAGGGCGGAAGTTTTTCAGAGGCGTTAACTGCGGCTTTGCGATCTCATG |
| SVEN0835 M3 F | CATGAGATCGCAAAGCCGCAGACAGCGAGCCGAGAGAACTTCCGCCCTAACGTGCAATG |
| SVEN0835 M3 R | CATTGCACGTTAGGGCGGAAGTTCTCTCGGCTCGCTGTCTGCGGCTTTGCGATCTCATG |
| SVEN1677 WT F | CTAGGCGAGAAGCACGCCATCATGTAACCTGCACGAAATTTCGCGGAGGGCCAACGTCG |
| SVEN1677 WT R | CGACGTTGGCCCTCCGCGAAATTTCGTGCAGGTTACATGATGGCGTGCTTCTCGCCTAG |
| SVEN1677 M1 F | CTAGGCGAGAAGCACGCCATCATTCAAGCTGCACAAATTTTCGCGGAGGGCCAACGTCG |
| SVEN1677 M1 R | CGACGTTGGCCCTCCGCGAAAATTTGTGCAGCTTGAATGATGGCGTGCTTCTCGCCTAG |
| SVEN1860 WT F | GTTTCATCAGTGTTTGACCACCGGGTCACGCCCTGGTA |

| | |
|---|---|
| | ACACCAGTCTGTGAGCCTGGT |
| SVEN1860 WT R | ACCAGGCTCACAGACTGGTGTTACCAGGGCGTGACCC GGTGGTCAAACACTGATGAAAC |
| SVEN1860 M1 F | GTTTCATCAGTGTTTGACCACCGGACCACGCCCTGACA ACACCAGTCTGTGAGCCTGGT |
| SVEN1860 M1 R | ACCAGGCTCACAGACTGGTGTTGTCAGGGCGTGGTCCG GTGGTCAAACACTGATGAAAC |
| SVEN1860 M2 F | GTTCCTTCAGTGTTCGGGCACCGGGTCACGCCCTGGTA ACACCAGTCTGTGAGCCTGGT |
| SVEN1860 M2 R | ACCAGGCTCACAGACTGGTGTTACCAGGGCGTGACCC GGTGCCCGAACACTGAAGGAAC |
| SVEN1860 M3 F | GTTCCTTCAGTGTTCGGGCACCGGACCACGCCCTGACA ACACCAGTCTGTGAGCCTGGT |
| SVEN1860 M3 R | ACCAGGCTCACAGACTGGTGTTGTCAGGGCGTGGTCCG GTGCCCGAACACTGAAGGAAC |
| SVEN1863 WT F | CCCACCCCGGTTAACTTCGACGAAACAATTGGGTCATG CTTGAGAAATCCCGTCTGCCT |
| SVEN1863 WT R | AGGCAGACGGGATTTCTCAAGCATGACCCAATTGTTTC GTCGAAGTTAACCGGGGTGGG |
| SVEN1863 M1 F | CCCACCCCGGTCAATTTCGACAGGATAATTGGGTCATGC TTGAGAAATCCCGTCTGCCT |
| SVEN1863 M1 R | AGGCAGACGGGATTTCTCAAGCATGACCCAATTATCCT GTCGAAATTGACCGGGGTGGG |
| SVEN1863 M2 F | CCCACCCCGGTTAACTTCGACGAAACAATTGGACCATG CTTGAAGGATCCCGTCTGCCT |
| SVEN1863 M2 R | AGGCAGACGGGATCCTTCAAGCATGGTCCAATTGTTTC GTCGAAGTTAACCGGGGTGGG |
| SVEN1863 M3 F | CCCACCCCGGTCAATTTCGACAGGATAATTGGACCATG CTTGAAGGATCCCGTCTGCCT |
| SVEN1863 M3 R | AGGCAGACGGGATCCTTCAAGCATGGTCCAATTATCCT GTCGAAATTGACCGGGGTGGG |
| SVEN1874 WT F | AGGAAAGCTGAGTAACACGGGGTTCACATTCGGGCAA CCGACGGGAAATCCCGTGTTGC |
| SVEN1874 WT R | GCAACACGGGATTTCCCGTCGGTTGCCCGAATGTGAAC CCCGTGTTACTCAGCTTTCCT |
| SVEN1874 M1 F | AGGAAAGCTGAACAAGACGGGGTCCAGATTCGGGCAA CCGACGGGAAATCCCGTGTTGC |
| SVEN1874 M1 R | GCAACACGGGATTTCCCGTCGGTTGCCCGAATCTGGAC CCCGTCTTGTTCAGCTTTCCT |
| SVEN1874 M2 F | AGGAAAGCTGAGTAACACGGGGTTCACATTCGGACAA GCGACGGAGGATCCCGTGTTGC |
| SVEN1874 M2 R | GCAACACGGGATCCTCCGTCGCTTGTCCGAATGTGAAC CCCGTGTTACTCAGCTTTCCT |
| SVEN1874 M3 F | AGGAAAGCTGAACAAGACGGGGTCCAGATTCGGACAA GCGACGGAGGATCCCGTGTTGC |

| | |
|---|---|
| SVEN1874 M3 R | GCAACACGGGATCCTCCGTCGCTTGTCCGAATCTGGAC CCCGTCTTGTTCAGCTTTCCT |
| SVEN2276 WT F | GGAGATCAGCGTTCCGTACGGGAAACAGTCGGGATGCC GCCGGGTAACACCGGCCGCGC |
| SVEN2276 WT R | GCGCGGCCGGTGTTACCCGGCGGCATCCCGACTGTTTC CCGTACGGAACGCTGATCTCC |
| SVEN2276 M1 F | GGAGATCAGCTCTCCGTACGGAAGATAGTCGGGATGCC GCCGGGTAACACCGGCCGCGC |
| SVEN2276 M1 R | GCGCGGCCGGTGTTACCCGGCGGCATCCCGACTATCTT CCGTACGGAGAGCTGATCTCC |
| SVEN2276 M2 F | GGAGATCAGCGTTCCGTACGGGAAACAGTCGGAACGG CGCCGGACAGCACCGGCCGCGC |
| SVEN2276 M2 R | GCGCGGCCGGTGCTGTCCGGCGCCGTTCCGACTGTTTC CCGTACGGAACGCTGATCTCC |
| SVEN2276 M3 F | GGAGATCAGCTCTCCGTACGGAAGATAGTCGGAACGGC GCCGGACAGCACCGGCCGCGC |
| SVEN2276 M3 R | GCGCGGCCGGTGCTGTCCGGCGCCGTTCCGACTATCTT CCGTACGGAGAGCTGATCTCC |
| SVEN2756 WT F | TACGGTGCAGGCCGACCACGGGGGCCCACGGGGGTGA CATCCATGTCTGGCATCAACAC |
| SVEN2756 WT R | GTGTTGATGCCAGACATGGATGTCACCCCCGTGGGCCC CCGTGGTCGGCCTGCACCGTA |
| SVEN2756 M1 F | TACGGTGCAGGCCGACCACGGGGGCCCACGGGGAGGC TATCCATGTCTGGCATCAACAC |
| SVEN2756 M1 R | GTGTTGATGCCAGACATGGATAGCCTCCCCGTGGGCCC CCGTGGTCGGCCTGCACCGTA |
| SVEN2756 M2 F | TACGGTGCAGGCCGACCACGGGGGCCCACGGGGGTGA CATCCATACTTGGCATCAACAC |
| SVEN2756 M2 R | GTGTTGATGCCAAGTATGGATGTCACCCCCGTGGGCCC CCGTGGTCGGCCTGCACCGTA |
| SVEN2756 M3 F | TACGGTGCAGGCCGACCACGGGGGCCCACGGGGAGGC TATCCATACTTGGCATCAACAC |
| SVEN2756 M3 R | GTGTTGATGCCAAGTATGGATAGCCTCCCCGTGGGCCCC CGTGGTCGGCCTGCACCGTA |
| SVEN3917 WT F | GCTCTCGCGCCCTGAGAGCTTTTGTTCATCCATCCGTAA CAACGGTCGGAAAACGCAAA |
| SVEN3917 WT R | TTTGCGTTTTCCGACCGTTGTTACGGATGGATGAACAA AAGCTCTCAGGGCGCGAGAGC |
| SVEN3917 M1 F | GCTCTCGCGCCCTGAGAGCTTTTGTCCCTCCATCCACA ACAACGGTCGGAAAACGCAAA |
| SVEN3917 M1 R | TTTGCGTTTTCCGACCGTTGTTGTGGATGGAGGGACAA AAGCTCTCAGGGCGCGAGAGC |
| SVEN3917 M2 F | GCTCTCGCGCCCTGAGAGCTTTTGTTCATGCCTCTGTAA CAACGGTCGGAAAACGCAAA |
| SVEN3917 M2 R | TTTGCGTTTTCCGACCGTTGTTACAGAGGCATGAACAA |

| | |
|---|---|
| | AAGCTCTCAGGGCGCGAGAGC |
| SVEN3917 M3 F | GCTCTCGCGCCCTGAGAGCTTTTGAGAGGCCATCCAGAACAACGGTCGGAAAACGCAAA |
| SVEN3917 M3 R | TTTGCGTTTTCCGACCGTTGTTCTGGATGGCCTCTCAAAAGCTCTCAGGGCGCGAGAGC |
| SVEN5279 WT F | CCGGCCGTTCACGGTCGCGTAACACGCCCCACCCCTTCGTCACGGCTCCGAAACATCGA |
| SVEN5279 WT R | TCGATGTTTCGGAGCCGTGACGAAGGGGTGGGGCGTGTTACGCGACCGTGAACGGCCGG |
| SVEN5279 M1 F | CCGGCCGTCCGCGGTCGCACAGCACGCCCCACCCCTTCGTCACGGCTCCGAAACATCGA |
| SVEN5279 M1 R | TCGATGTTTCGGAGCCGTGACGAAGGGGTGGGGCGTGCTGTGCGACCGCGGACGGCCGG |
| SVEN5279 M2 F | CCGGCCGTTCACGGTCGCGTAACACGCCCCACCCCTTCACCGCGGCTCCAGGATATCGA |
| SVEN5279 M2 R | TCGATATCCTGGAGCCGCGGTGAAGGGGTGGGGCGTGTTACGCGACCGTGAACGGCCGG |
| SVEN5279 M3 F | CCGGCCGTCCGCGGTCGCACAGCACGCCCCACCCCTTCACCGCGGCTCCAGGATATCGA |
| SVEN5279 M3 R | TCGATATCCTGGAGCCGCGGTGAAGGGGTGGGGCGTGCTGTGCGACCGCGGACGGCCGG |
| SVEN6462 WT F | CGGCCCGCGGCGGGAACGTTCGGGCCGGTGCGTCATACTGAACGACCATTCAGTACGGC |
| SVEN6462 WT R | GCGGTGAGCGGTCGGAGCCGGTACCGGTCACAGGACCTCCTGATCTCTGGGGGTCGTTC |
| **For ChIP-qPCR analysis** | |
| hrdB(sven5499) chip-qPCR F | GCCGAGGAAGGAATACAGCA |
| hrdB(sven5499) chip-qPCR R | AGGTCCTGGAGCATCTGGC |
| sven0835 chip-qPCR F | GCAGTTAACGAGCCGGAAAA |
| sven0835 chip-qPCR R | ACACCTTGTTGAACTGCGGA |
| sven1677 chip-qPCR F | GAAGCACGCCATCATGTAACC |
| sven1677 chip-qPCR R | ATATGCAGATGCCGAGGGA |
| sven1860 chip-qPCR F | ACGTTTCATCAGTGTTTGACCA |
| sven1860 chip-qPCR R | CGGTGAGTGAAACCAGGCT |
| sven1863 chip-qPCR F | CACCCACCCCGGTTAACTT |
| sven1863 chip-qPCR R | CTGGACCCGACCATAGGCAG |
| sven1874 chip-qPCR F | CCGCGAAGATCAGGAAAGC |
| sven1874 chip-qPCR R | GTCGCAGCTTCGCAACAC |
| sven2276 chip-qPCR F | CGGAGATCAGCGTTCCGTA |
| sven2276 chip-qPCR R | GTCTTCATACCGGGAAGCGT |
| sven2756 chip-qPCR F | TGACATCCATGTCTGGCATCA |
| sven2756 chip-qPCR R | GAACGCGTCCCTTCATATCG |
| sven3917 chip-qPCR F | ACTGCCGGCTCTCGCGCCCTGA |
| sven3917 chip-qPCR R | AGCCCCCGGGTCCTTTTGCGTT |

| | |
|---|---|
| sven5279 chip-qPCR F | CGTTCACGGTCGCGTAACAC |
| sven5279 chip-qPCR R | GTTTCCGTCGATGCCGCTC |
| sven6462 chip-qPCR F | CCTGATCTCTGGGGGTCGTT |
| sven6462 chip-qPCR R | TCGGCCGTACTGAATGGTC |
| **For Real-time PCR analysis** | |
| hrdB(sven5499) realtime F | CCAGATTCCGCCAACCCA |
| hrdB(sven5499) realtime R | CTTCGTCACGGTCGTCCTG |
| sven0835 realtime F | GAGCCCGCACGAACAGGAA |
| sven0835 realtime R | CAGGATGTGGGAGGTGAGGAG |
| sven1677 realtime F | TTCGTCGCCCACAACGGTGAGAT |
| sven1677 realtime R | GGTGCAGATGGGGAAGATCCGCT |
| sven1860 realtime F | ATGCCGGCCGCATCGACGCA |
| sven1860 realtime R | TCCAGGACGGAGTTCCAGGC |
| sven1863 realtime F | TCTACGACGAGACGGGCTACG |
| sven1863 realtime R | CAGGCGGTGGTACGAGTTCA |
| sven1874 realtime F | GGACCGTGTTCTCAAGCC |
| sven1874 realtime R | GTTGGACTCGTGCGGCGT |
| sven2277 realtime F | CCAGGATCGTGGTCCTCTGCGAG |
| sven2277 realtime R | GGGTCTTGCCCGAGAAGTACGAGGT |
| sven2756-wt realtime F | AGACTGATACGGAAATGGGA |
| sven2756-wt realtime R | TCTCGGCCAGTGCGGTGTCGT |
| sven3917-wt realtime F | GACCCGGACGTTGTGCAG |
| sven3917-wt realtime R | GCCCTGCTGCTCCTGAC |
| sven2756 realtime F | GTGCTGCGGGGTGAAGGGTT |
| sven2756 realtime R | CGTGTCGCTCTTGGCCGTGAG |
| sven3917 realtime F | TCGGGGGTGACGAAGCGG |
| sven3917 realtime R | CCAGGAGGTGTGGGGGTA |
| sven5279 realtime F | GCTGGTCCTCCGACTACG |
| sven5279 realtime R | GGTGAGGATGGCGAACAT |
| sven5280 realtime F | GGTCCCCAAGATCCGCAT |
| sven5280 realtime R | ACACCTTGCCGTCTCCGA |
| sven5281 realtime F | TCCTCCTGCTCCACGACG |
| sven5281 realtime R | GTGCGGACGGAGTGGTCG |
| sven6462 realtime F | CTGCGGATGCGGGAGGGCT |
| sven6462 realtime R | GCCCGAGGGTCGTAGTGC |

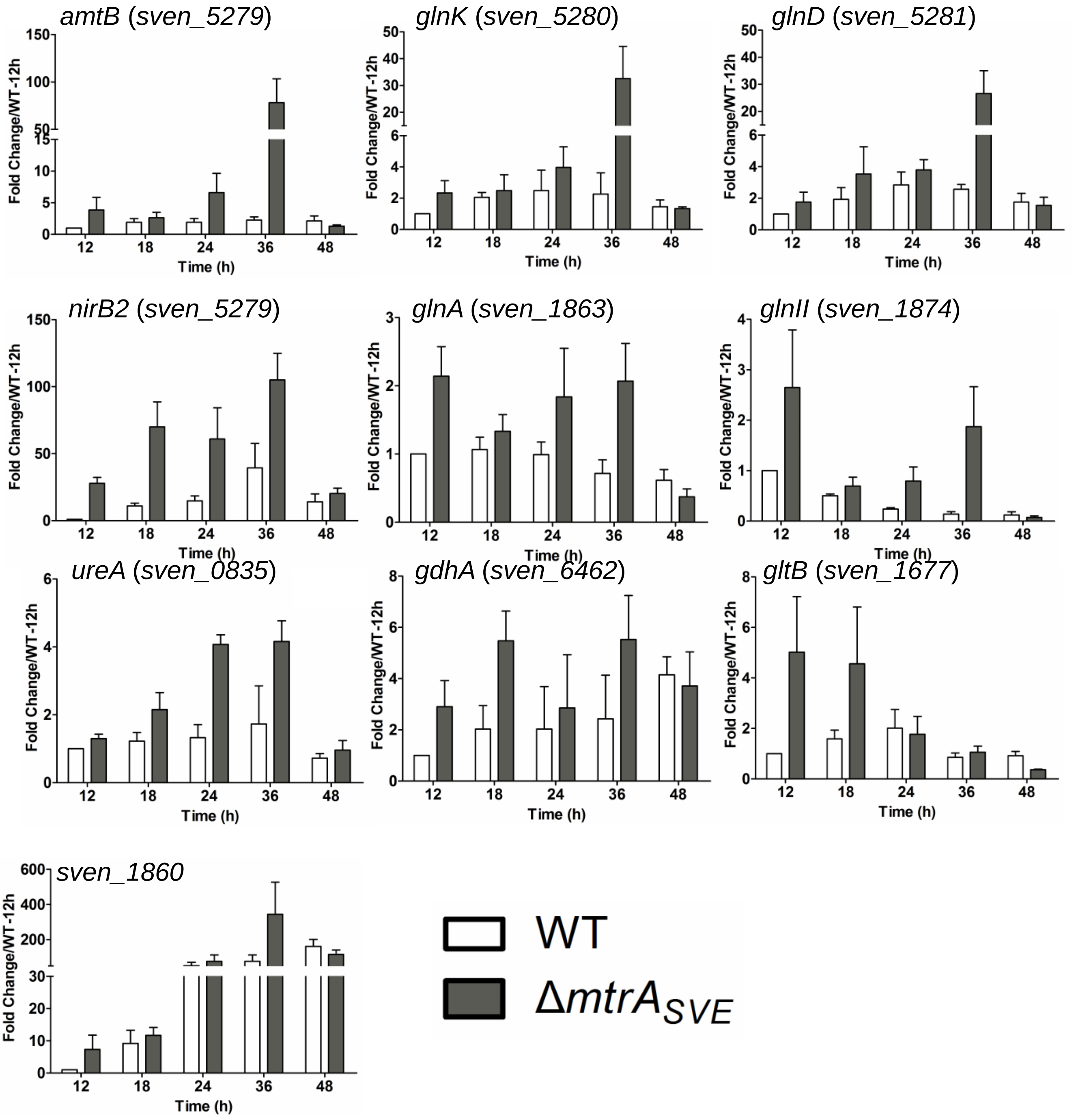


Figure S1. Transcriptional analysis of nitrogen metabolism genes in the Δ*mtrA*$_{SVE}$ mutant by real-time PCR. *Streptomyces* strains were cultured on solid YBP, and RNA samples from the wild-type strain 10712 (WT) and Δ*mtrA*$_{SVE}$ were isolated at the indicated times. Expression of *hrdB,* encoding the major sigma factor, was used as an internal control. For each gene, the expression level in WT at the first time point was arbitrarily set to one. The y-axis shows the fold change in expression in WT and Δ*mtrA*$_{SVE}$ over the expression levels in WT at the first time point. Results are the means (±SD) of triplet biological experiments.

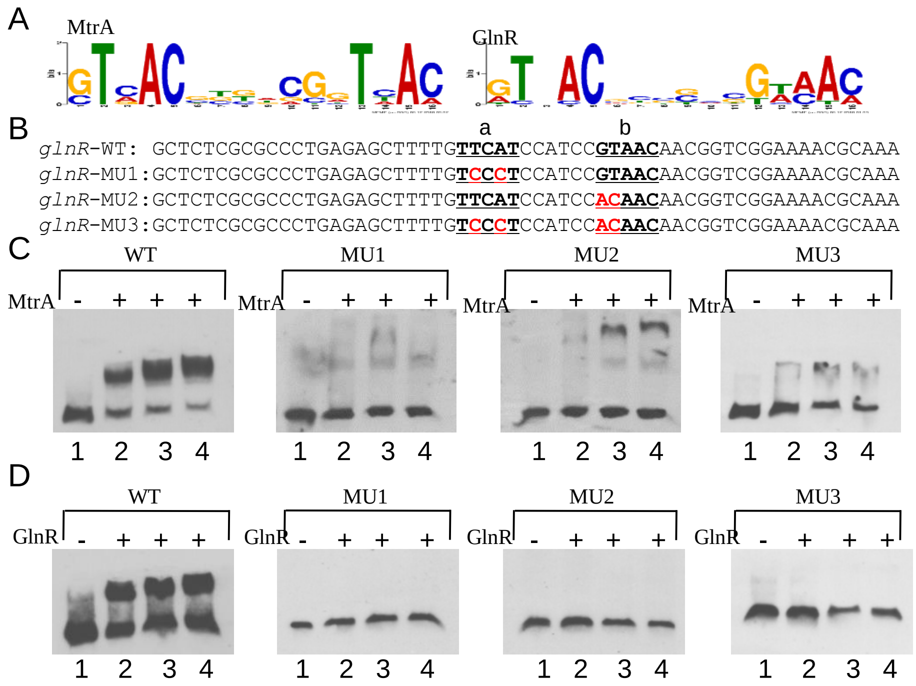


Figure S2. Mutational analyses of the MtrA site/GlnR box in the *glnR* promoter. (A) Consensus binding motifs for MtrA and GlnR. (B) The sequences of the WT and mutagenized probes for the MtrA site/GlnR box of the *glnR* promoter. The core nucleotides of the MtrA site/GlnR box are underlined, and mutagenized bases are shown in red. Probes were used in EMSAs with (C) MtrA or (D) GlnR. Reactions were carried out with the addition of no MtrA (lanes 1), 1.18 μM MtrA (lanes 2), 4.74 μM MtrA (lanes 3), or 8.29 μM MtrA (lanes 4) for panel C; or no GlnR (lanes 1), 0.45 μM GlnR (lanes 2), 1.78 μg GlnR (lanes 3), or 3.11 μM GlnR (lanes 4) for panel D.

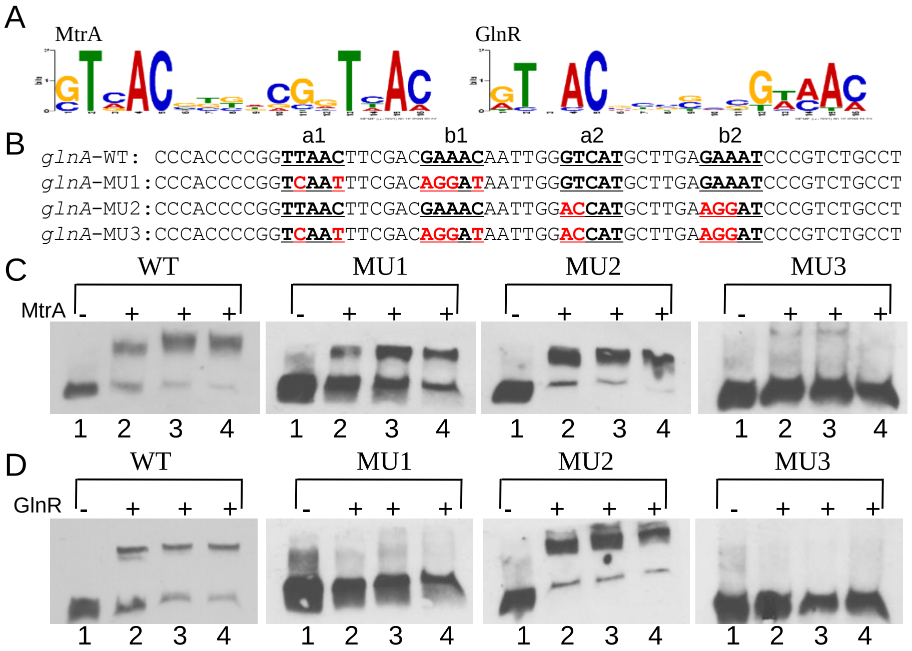


Figure S3. Mutational analyses of the MtrA sites/GlnR boxes in the *glnA* promoter. (A) Consensus binding motifs for MtrA and GlnR. (B) The sequences of the WT and mutagenized probes for MtrA sites/GlnR boxes of the *glnA* promoter. The core nucleotides of the MtrA sites/GlnR boxes are underlined, and mutagenized bases are shown in red. Probes were used in EMSAs with (C) MtrA or (D) GlnR. Reactions were carried out with the addition of no MtrA (lanes 1), 1.18 μM MtrA (lanes 2), 4.74 μM MtrA (lanes 3), or 8.29 μM MtrA (lanes 4) for panel C; or no GlnR (lanes 1), 0.45 μM GlnR (lanes 2), 1.78 μg GlnR (lanes 3), or 3.11 μM GlnR (lanes 4) for panel D.

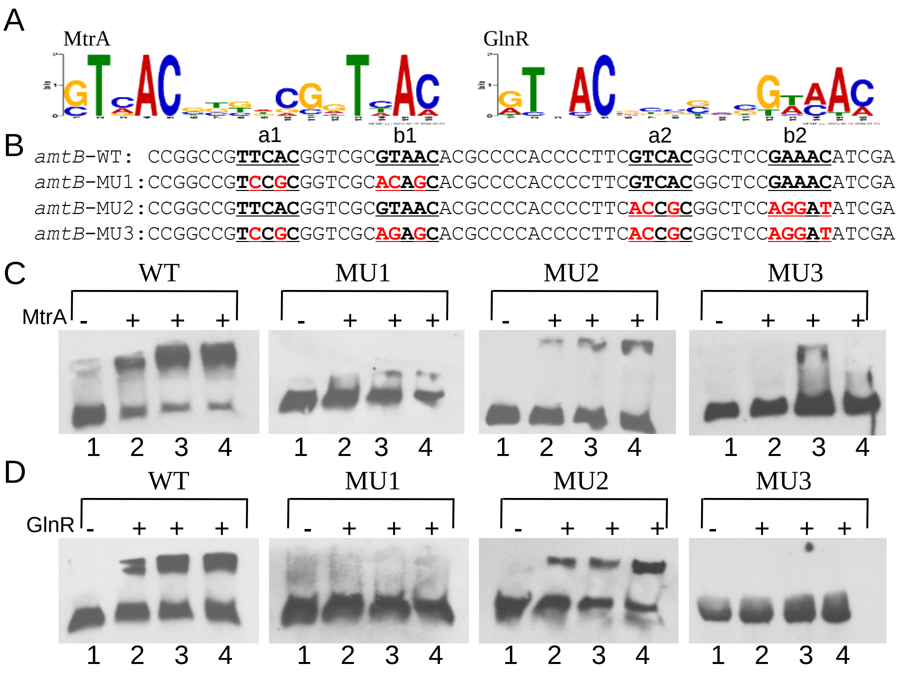


Figure S4. Mutational analyses of the MtrA/GlnR boxes in the *amtB* promoter. (A) Consensus binding motifs for MtrA and GlnR. (B) The sequences of the WT and mutagenized probes for MtrA sites/GlnR boxes of the *amtB* promoter. The core nucleotides of the MtrA sites/GlnR boxes are underlined, and mutagenized bases are shown in red. Probes were used in EMSAs with (C) MtrA or (D) GlnR. Reactions were carried out with the addition of no MtrA (lanes 1), 1.18 μM MtrA (lanes 2), 4.74 μM MtrA (lanes 3), or 8.29 μM MtrA (lanes 4) for panel C; or no GlnR (lanes 1), 0.45 μM GlnR (lanes 2), 1.78 μg GlnR (lanes 3), or 3.11 μM GlnR (lanes 4) for panel D.

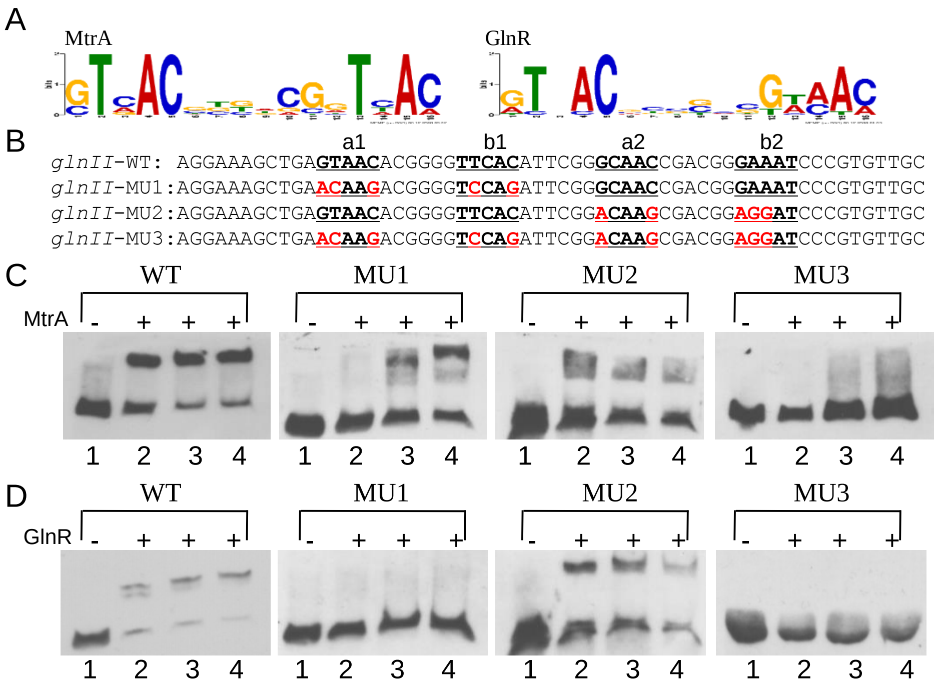


Figure S5. Mutational analyses of the MtrA sites/GlnR boxes in the *glnII* promoter. (A) Consensus binding motifs for MtrA and GlnR. (B) The sequences of the WT and mutagenized probes for MtrA sites/GlnR boxes of the *glnII* promoter. The core nucleotides of the MtrA site/GlnR boxes are underlined, and mutagenized bases are shown in red. Probes were used in EMSAs with (C) MtrA or (D) GlnR. Reactions were carried out with the addition of no MtrA (lanes 1), 1.18 μM MtrA (lanes 2), 4.74 μM MtrA (lanes 3), or 8.29 μM MtrA (lanes 4) for panel C; or no GlnR (lanes 1), 0.45 μM GlnR (lanes 2), 1.78 μg GlnR (lanes 3), or 3.11 μM GlnR (lanes 4) for panel D.

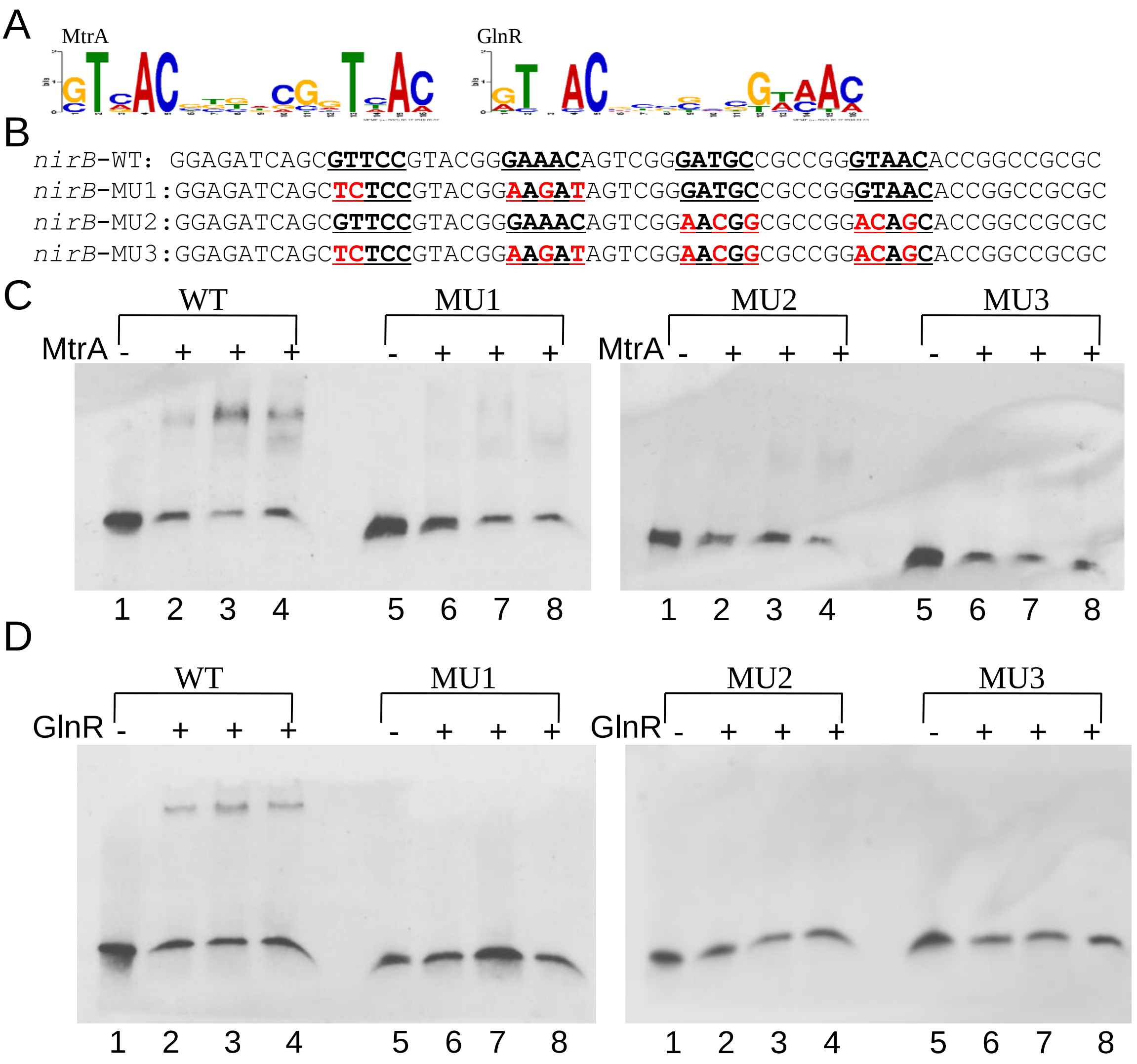


Figure S6. Mutational analyses of the MtrA sites/GlnR boxes in the *nirB* promoter. (A) Consensus binding motifs for MtrA and GlnR. (B) The sequences of the WT and mutagenized probes for MtrA sites/GlnR boxes of the *nirB* promoter. The core nucleotides of the MtrA sites/GlnR boxes are underlined, and mutagenized bases are shown in red. Probes were used in EMSAs with (C) MtrA or (D) GlnR. Reactions were carried out with the addition of no MtrA (lanes 1), 1.18 μM MtrA (lanes 2), 4.74 μM MtrA (lanes 3), or 8.29 μM MtrA (lanes 4) for panel C; or no GlnR (lanes 1), 0.45 μM GlnR (lanes 2), 1.78 μg GlnR (lanes 3), or 3.11 μM GlnR (lanes 4) for panel D.

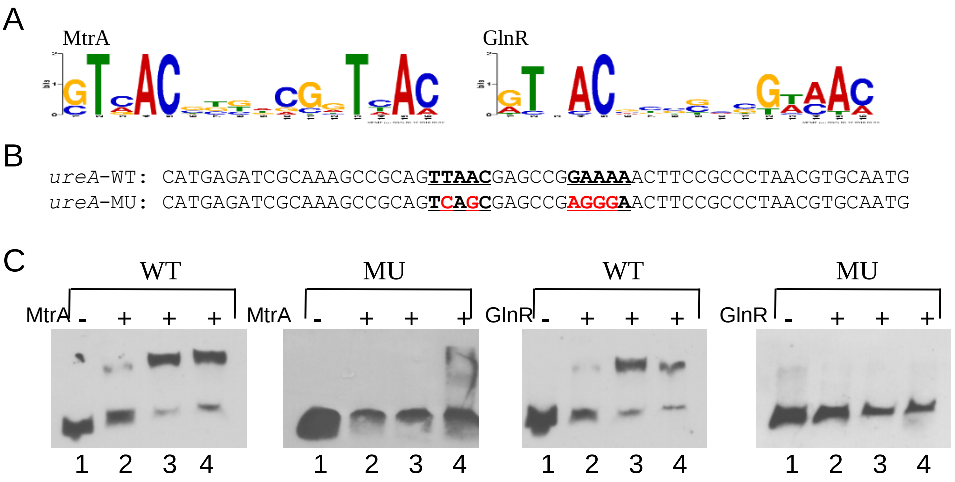


Figure S7. Mutational analyses of the MtrA site/GlnR box in the *ureA* promoter. (A) Consensus binding motifs for MtrA and GlnR. (B) The sequences of the WT and mutagenized probes for the MtrA site/GlnR box of the *ureA* promoter. The core nucleotides of the MtrA site/GlnR box are underlined, and mutagenized bases are shown in red. Probes were used in EMSAs with (C) MtrA or GlnR. Reactions were carried out with the addition of no MtrA (lanes 1), 1.18 μM MtrA (lanes 2), 4.74 μM MtrA (lanes 3), or 8.29 μM MtrA (lanes 4); or no GlnR (lanes 1), 0.45 μM GlnR (lanes 2), 1.78 μg GlnR (lanes 3), or 3.11 μM GlnR (lanes 4).

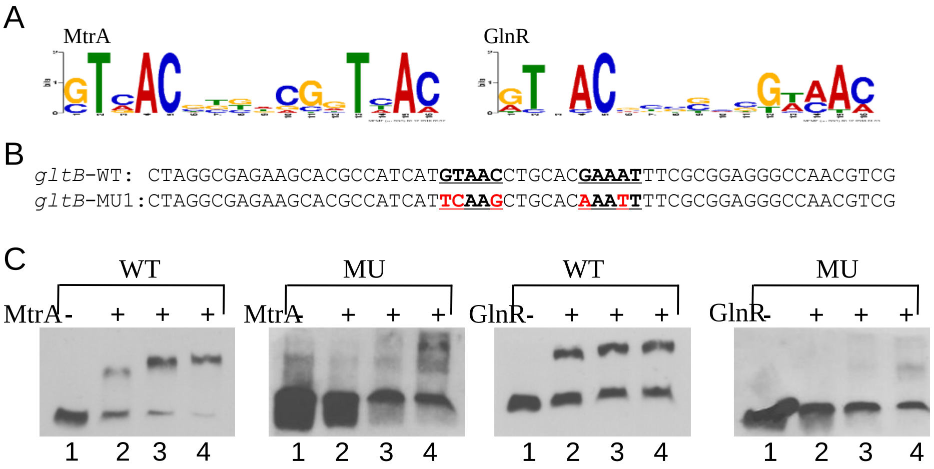


Figure S8. Mutational analyses of the MtrA site/GlnR box in the *gltB* promoter. (A) Consensus binding motifs for MtrA and GlnR. (B) The sequences of the WT and mutagenized probes for MtrA site/GlnR box of the *gltB* promoter. The core nucleotides of the MtrA site/GlnR box are underlined, and mutagenized bases are shown in red. Probes were used in EMSAs with (C) MtrA or GlnR. Reactions were carried out with the addition of no MtrA (lanes 1), 1.18 μM MtrA (lanes 2), 4.74 μM MtrA (lanes 3), or 8.29 μM MtrA (lanes 4); or no GlnR (lanes 1), 0.45 μM GlnR (lanes 2), 1.78 μg GlnR (lanes 3), or 3.11 μM GlnR (lanes 4).

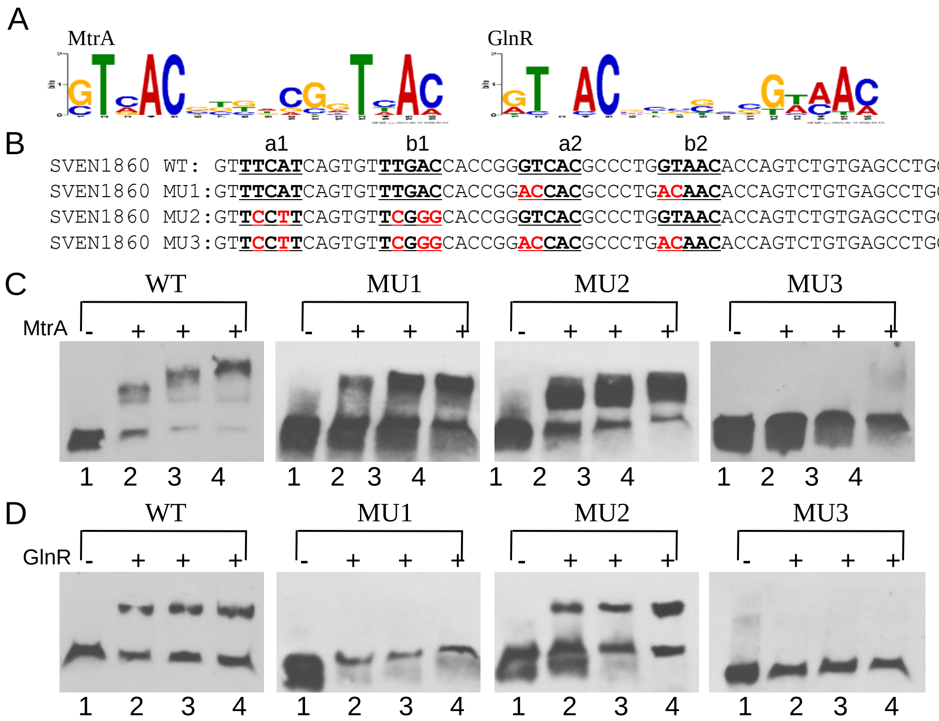


Figure S9. Mutational analyses of the MtrA sites/GlnR boxes in the *sven_1860* promoter. (A) Consensus binding motifs for MtrA and GlnR. (B) The sequences of the WT and mutagenized probes for MtrA sites/GlnR boxes of the *sven_1860* promoter. The core nucleotides of the MtrA sites/GlnR boxes are underlined, and mutagenized bases are shown in red. Probes were used in EMSAs with (C) MtrA or (D) GlnR. Reactions were carried out with the addition of no MtrA (lanes 1), 1.18 μM MtrA (lanes 2), 4.74 μM MtrA (lanes 3), or 8.29 μM MtrA (lanes 4) for panel C; or no GlnR (lanes 1), 0.45 μM GlnR (lanes 2), 1.78 μg GlnR (lanes 3), or 3.11 μM GlnR (lanes 4) for panel D.

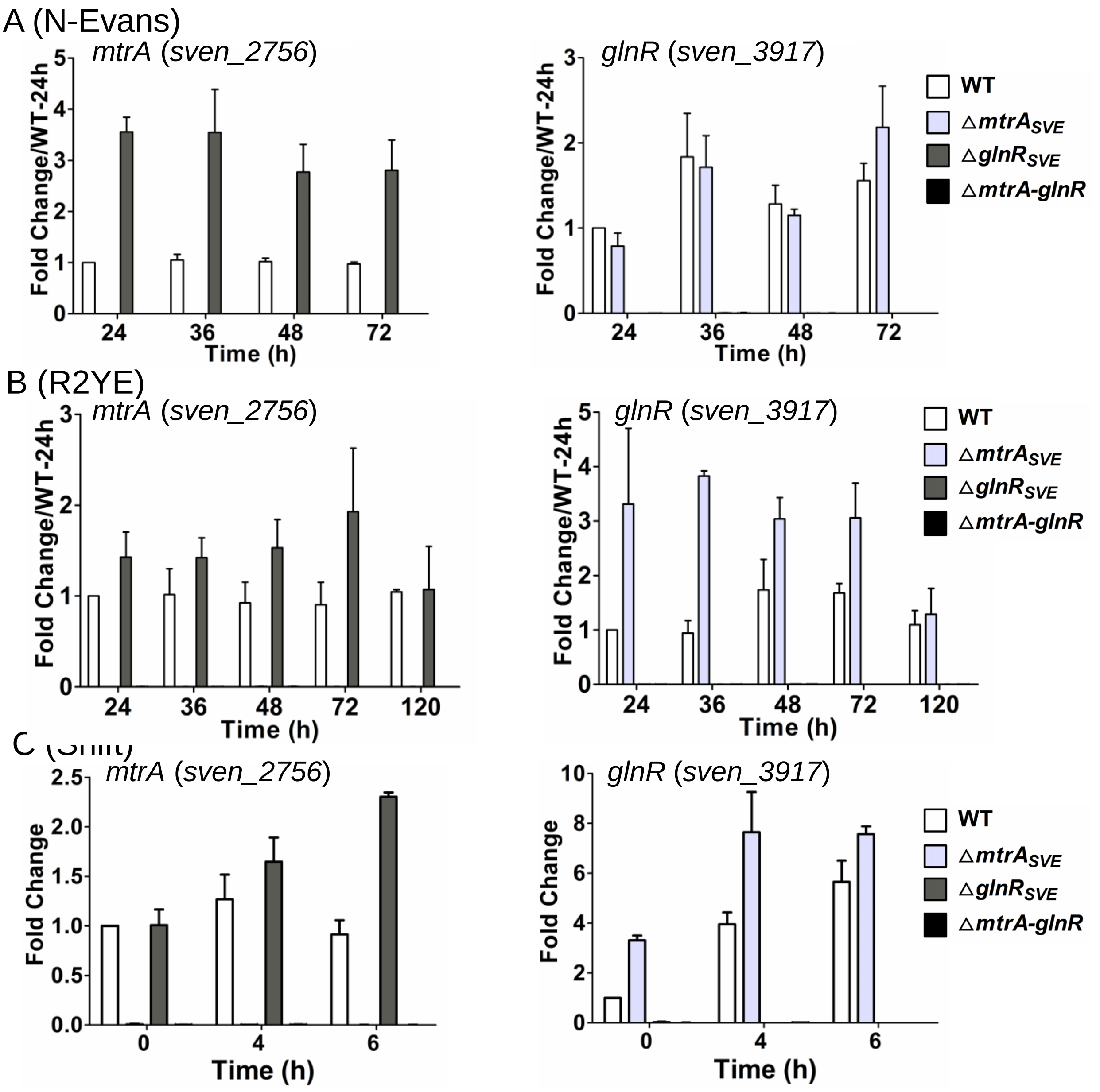


Figure S10. Transcriptional analysis of *mtrA* and *glnR* in Δ*mtrA*$_{SVE}$, Δ*glnR*$_{SVE}$, and Δ*mtrA-glnR* mutants by real-time PCR. *Streptomyces* strains were cultured on solid R2YE (A), N-Evans (B), or shifted from liquid YBP to liquid N-R2YE (C), and RNA samples from WT, Δ*mtrA*$_{SVE}$, Δ*glnR*$_{SVE}$, and Δ*mtrA-glnR* were isolated at the indicated times. Expression of *hrdB,* encoding the major sigma factor, was used as an internal control. For each gene, the expression level in WT at the first time point was arbitrarily set to one. The y-axis shows the fold change in expression in WT, Δ*mtrA*$_{SVE}$, Δ*glnR*$_{SVE}$, and Δ*mtrA-glnR* over the expression levels in WT at the first time point. Primer sets that target the deleted sequences in Δ*mtrA*$_{SVE}$ and Δ*glnR*$_{SVE}$ were used. Results are the means (±SD) of triplet biological experiments.

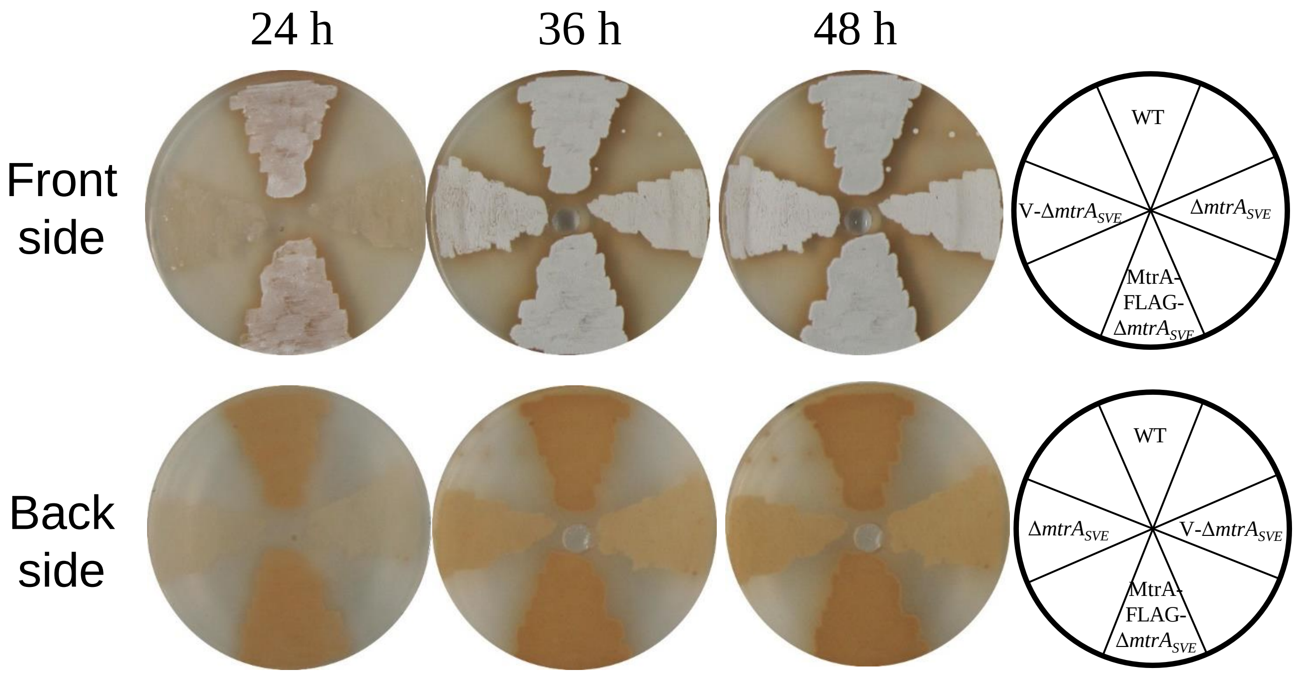


Figure S11. Restoration of pigment production and aerial mycelium formation with an MtrA-FLAG construct. *Streptomyces* strains were incubated on MS-G medium for the indicated times. The white patches are aerial mycelia. WT, ATCC 10172; $\Delta mtrA_{SVE}$, *mtrA* deletion mutant; MtrA-FLAG-$\Delta mtrA_{SVE}$, $\Delta mtrA_{SVE}$ complemented by pMtrA-FLAG; V-$\Delta mtrA_{SVE}$, $\Delta mtrA_{SVE}$ containing pMS82.

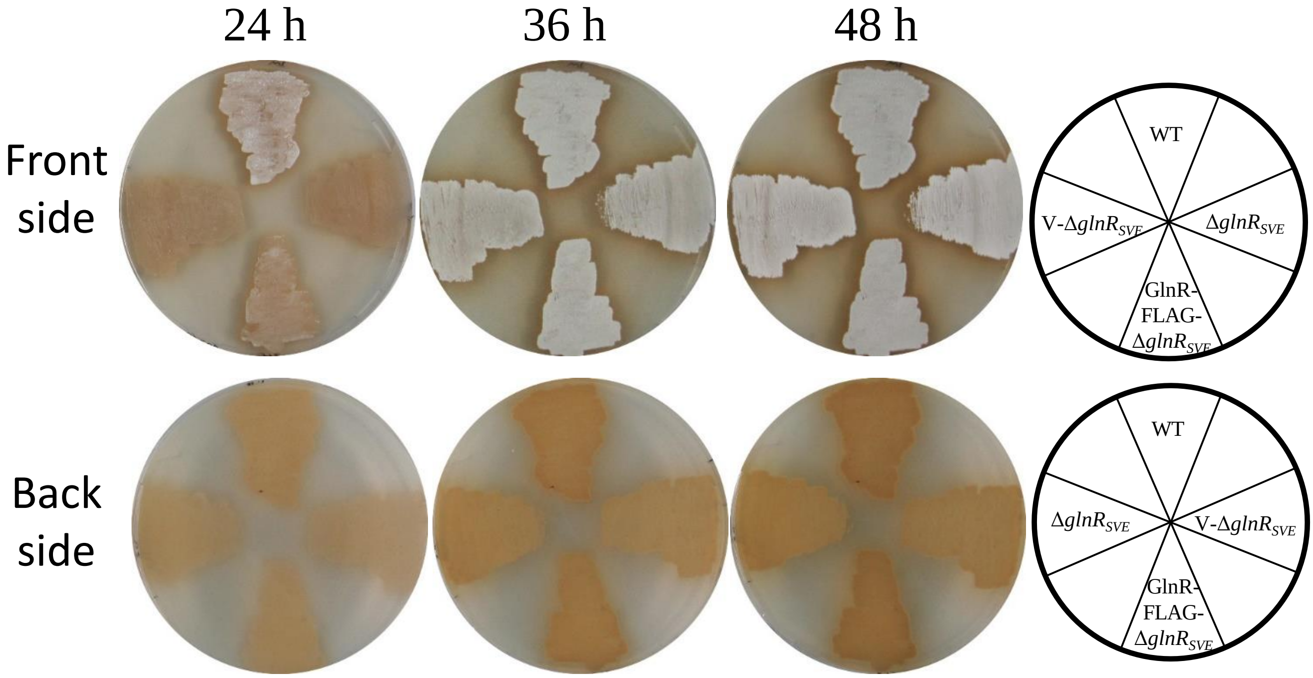


Figure S12. Restoration of pigment production and aerial mycelium formation with an GlnR-FLAG construct. *Streptomyces* strains were incubated on MS-G medium for the indicated times. The white patches are aerial mycelia. WT, ATCC 10172; Δ$glnR_{SVE}$, *glnR* deletion mutant; GlnR-FLAG-Δ$glnR_{SVE}$, Δ$glnR_{SVE}$ complemented by pGlnR-FLAG; V-Δ$glnR_{SVE}$, Δ$glnR_{SVE}$ containing pMS82.

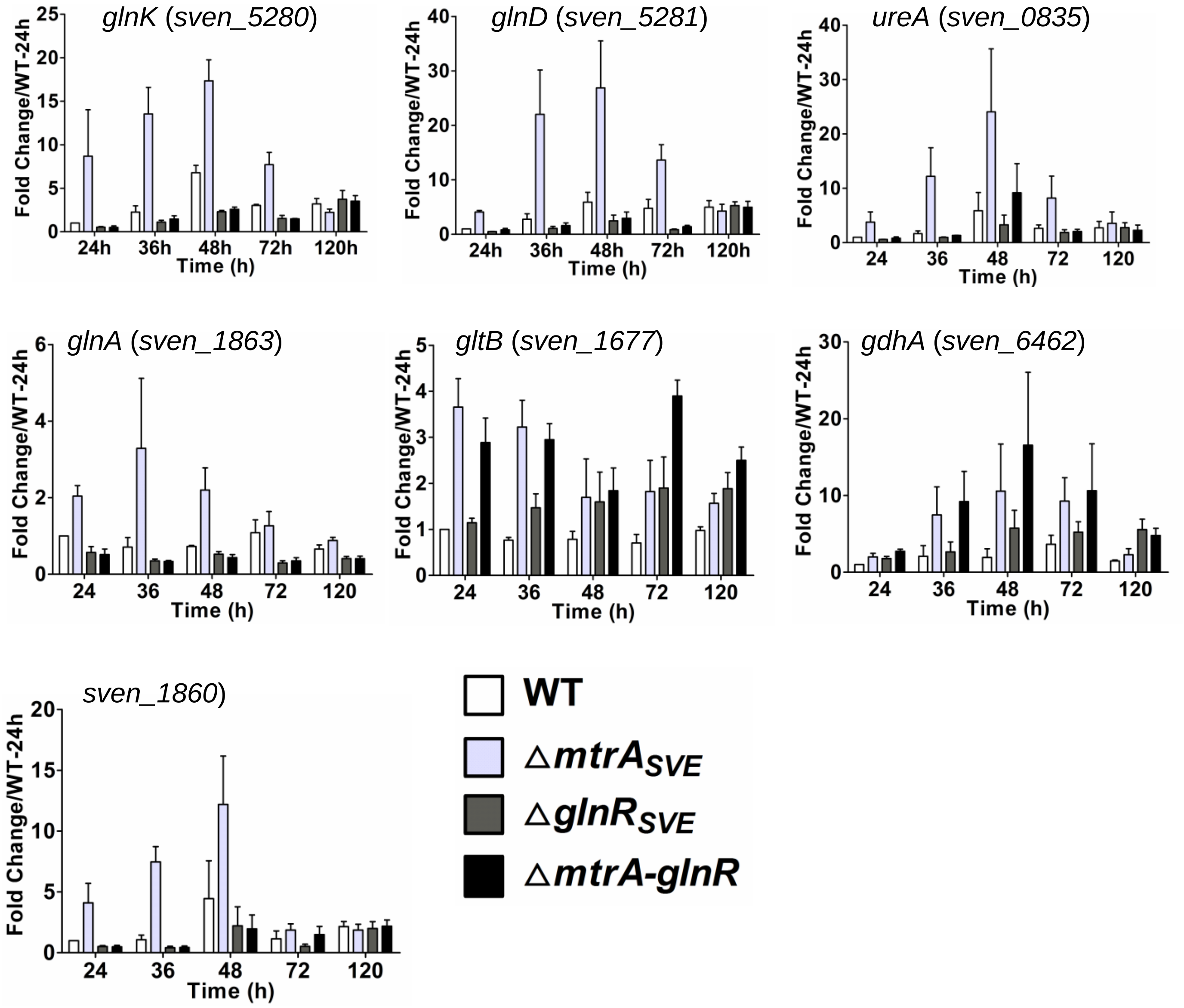


Figure S13. Transcriptional analysis by real-time PCR of nitrogen metabolism genes in Δ*mtrA*$_{SVE}$, Δ*glnR*$_{SVE}$, and Δ*mtrA-glnR* mutants grown on solid R2YE. RNA samples from WT, Δ*mtrA*$_{SVE}$, Δ*glnR*$_{SVE}$, and Δ*mtrA-glnR* were isolated at the indicated times. Expression of *hrdB,* encoding the major sigma factor, was used as an internal control. For each gene, the expression level in WT at the first time point was arbitrarily set to one. The y-axis shows the fold change in expression in WT, Δ*mtrA*, Δ*glnR*, and Δ*mtrA-glnR* over the expression levels in WT at the first time point. Results are the means (±SD) of triplet biological experiments.

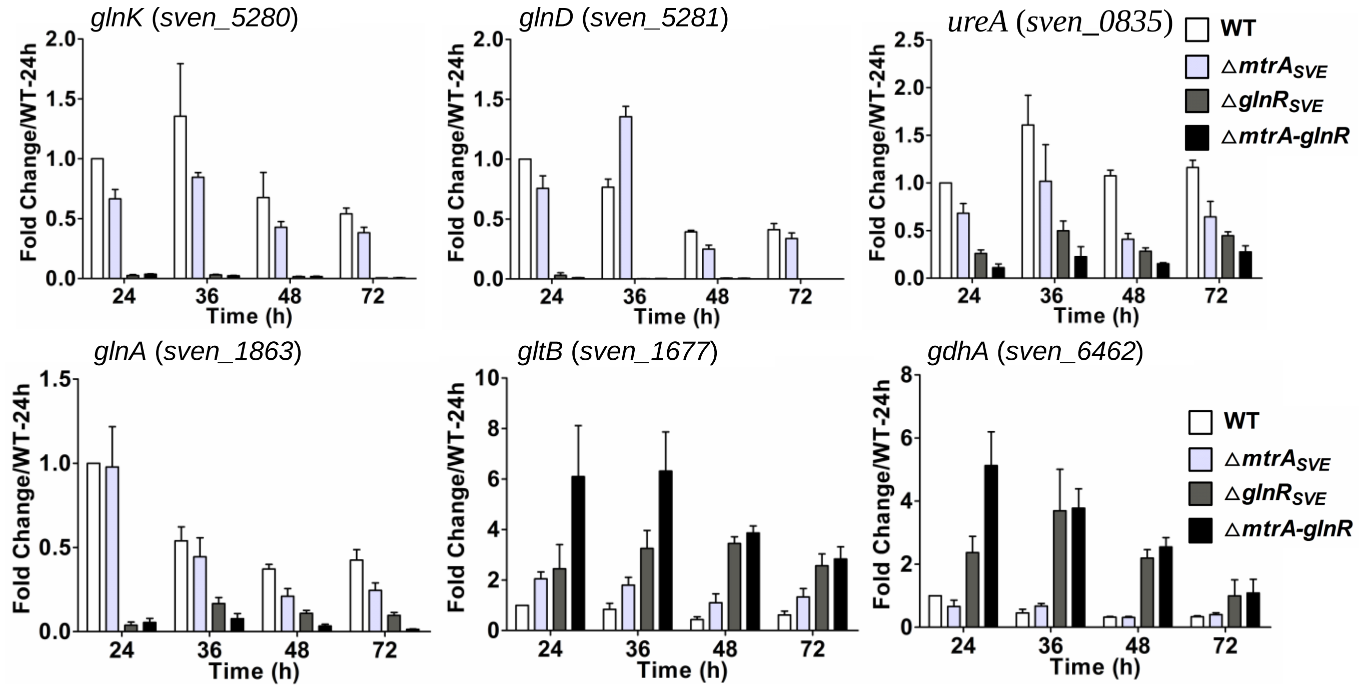


Figure S14. Transcriptional analysis by real-time PCR of nitrogen metabolism genes in Δ*mtrA*$_{SVE}$, Δ*glnR*$_{SVE}$, and Δ*mtrA-glnR* mutants grown on solid N-Evans. RNA samples from WT, Δ*mtrA*$_{SVE}$, Δ*glnR*$_{SVE}$, and Δ*mtrA-glnR* were isolated at the indicated times, and gene expression was determined as for Figure S5. Results are the means (±SD) of triplet biological experiments.

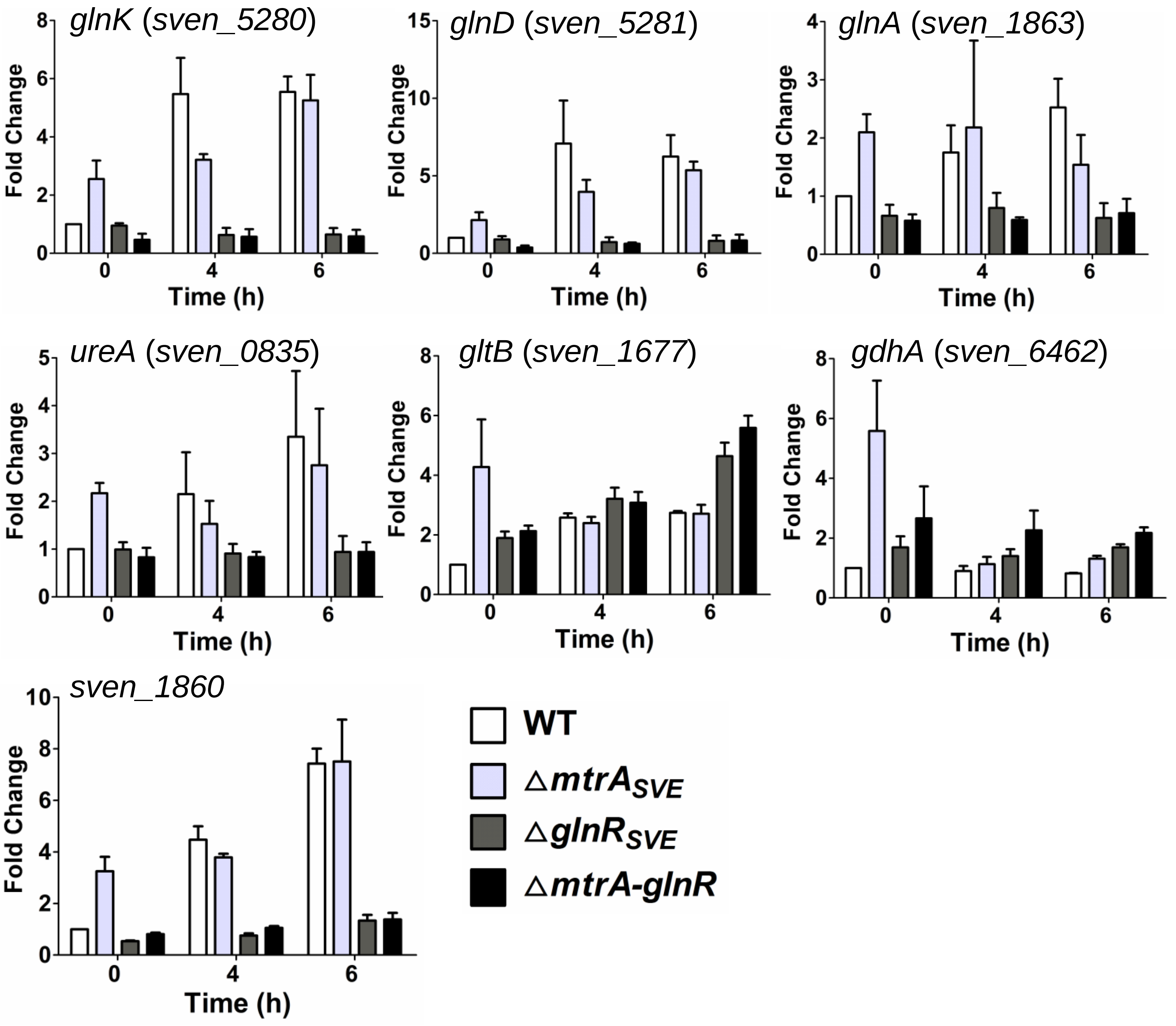


Figure S15. Transcriptional analysis by real-time PCR of nitrogen metabolism genes in Δ*mtrA*$_{SVE}$, Δ*glnR*$_{SVE}$, and Δ*mtrA-glnR* mutants under nutrient-shift. *Streptomyces* strains were cultured in liquid YBP and then transferred to liquid N-Evans. RNA samples from WT, Δ*mtrA*$_{SVE}$, Δ*glnR*$_{SVE}$, and Δ*mtrA-glnR* were isolated at the indicated times, and gene expression was determined as for Figure S5. Results are the means (±SD) of triplet biological experiments.

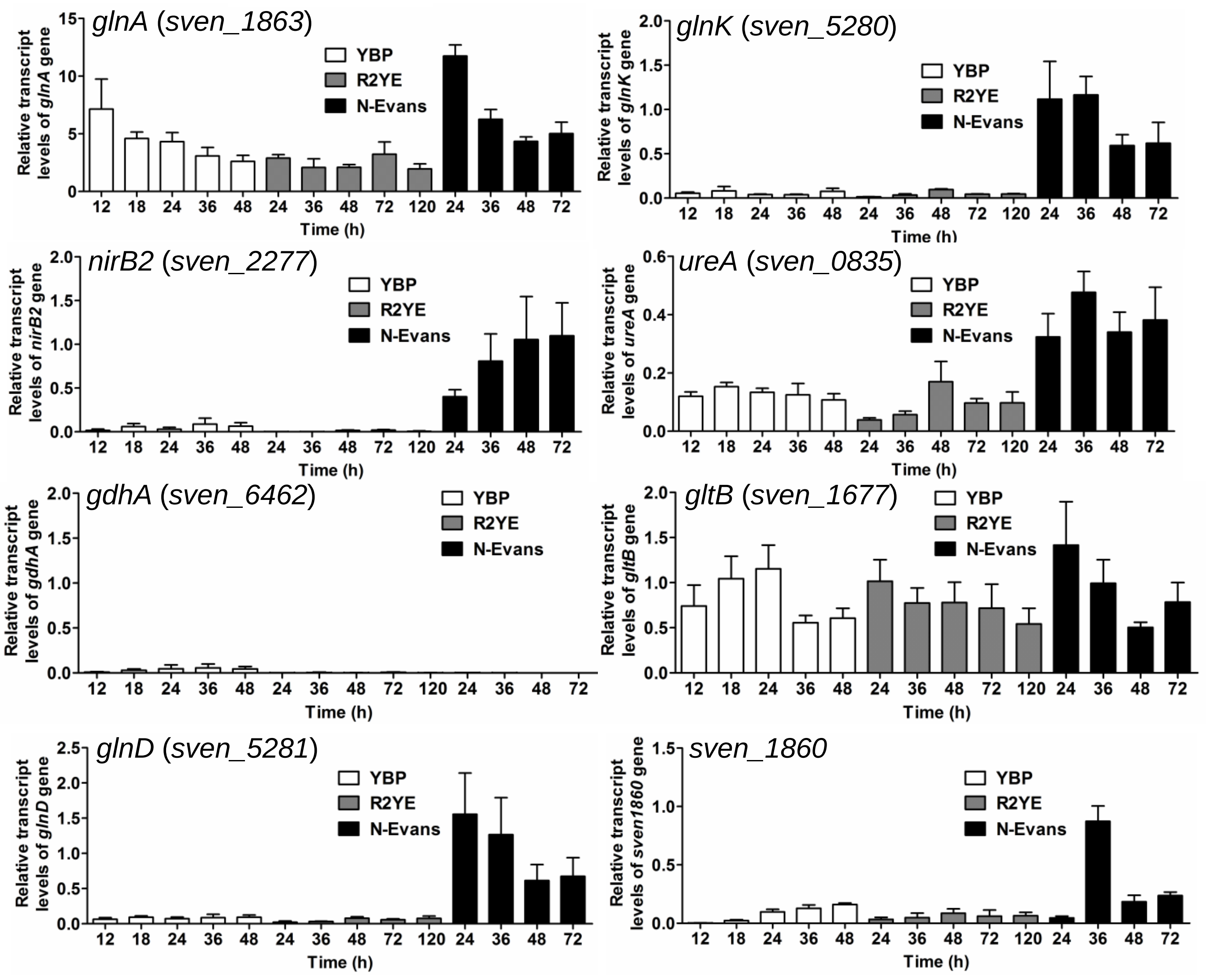


Figure S16. Transcriptional analysis by real-time PCR of nitrogen metabolism genes in the wild-type strain 10712 grown on different media. The wild-type strain was grown on YBP, R2YE, and N-Evans. For each gene, the expression level of *hrdB* at each time point was arbitrarily set to one. The y-axis shows the fold change in expression of each gene over the expression level of *hrdB* for each time point. Results are the means (±SD) of triplet biological experiments.

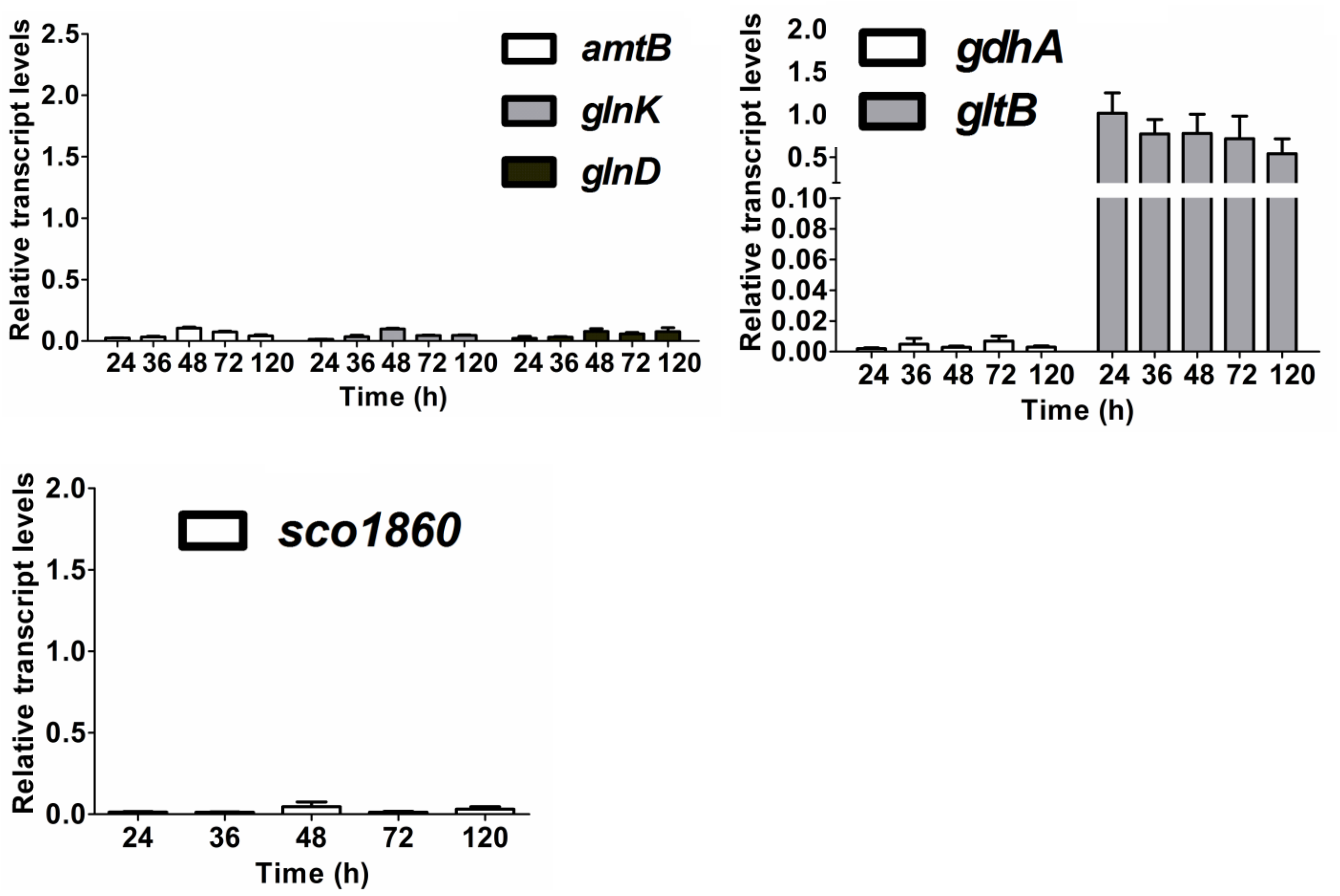


Figure S17. Transcriptional analysis by real-time PCR of nitrogen metabolism genes in the wild-type strain 10712 grown on R2YE. For each gene, the expression level of *hrdB* at each time point was arbitrarily set to one. The y-axis shows the fold change in expression of each gene over the expression level of *hrdB* for each time point. Results are the means (±SD) of triplet biological experiments.

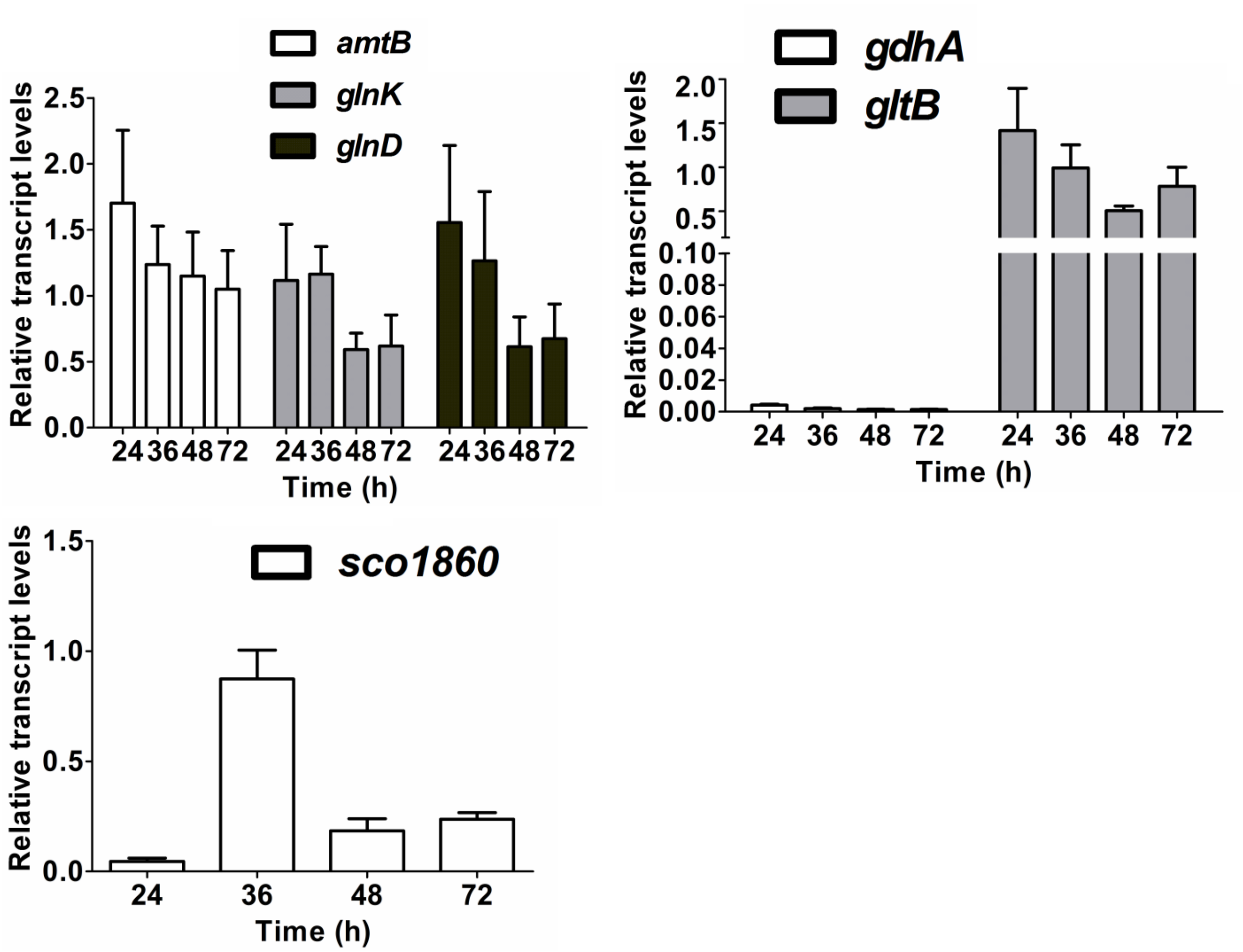


Figure S18. Transcriptional analysis by real-time PCR of nitrogen metabolism genes in the wild-type strain 10712 grown on N-Evans. For each gene, the expression level of *hrdB* at each time point was arbitrarily set to one. The y-axis shows the fold change in expression of each gene over the expression level of *hrdB* for each time point. Results are the means (±SD) of triplet biological experiments.

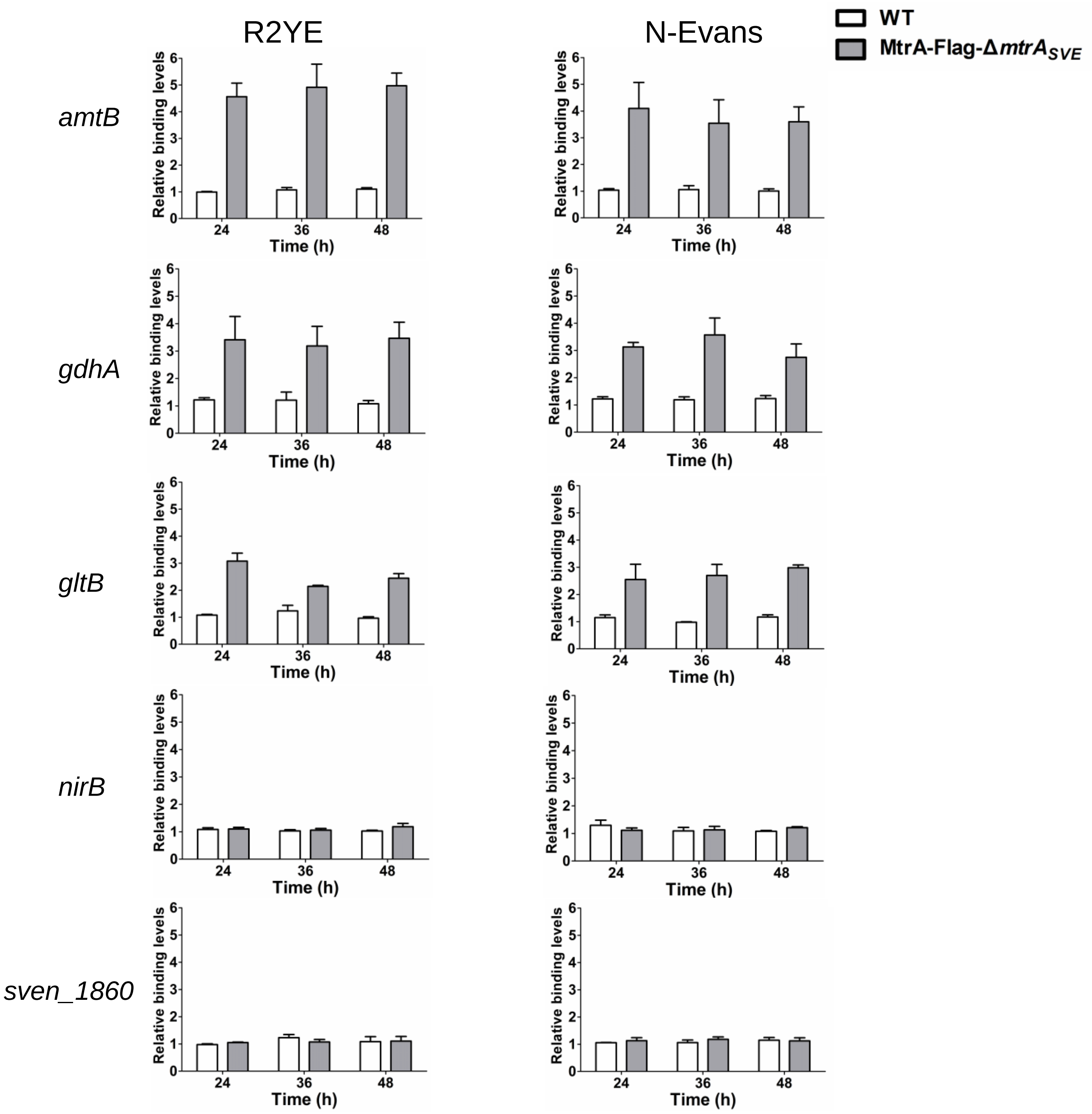


Figure S19. ChIP-qPCR analysis of the binding of MtrA to the promoters of nitrogen metabolism genes from cultures grown on R2YE or N-Evans. Analysis was performed using strains MtrA-FLAG-Δ*mtrA*$_{SVE}$ and the wild-type 10712 (WT) grown on R2YE or N-Evans for the indicated times. The y-axis shows the binding levels of MtrA-FLAG in MtrA-FLAG-Δ*mtrA*$_{SVE}$ and WT relative to background levels, which was determined by recovery of target sequences. Results are the means (±SD) of triplet biological experiments.

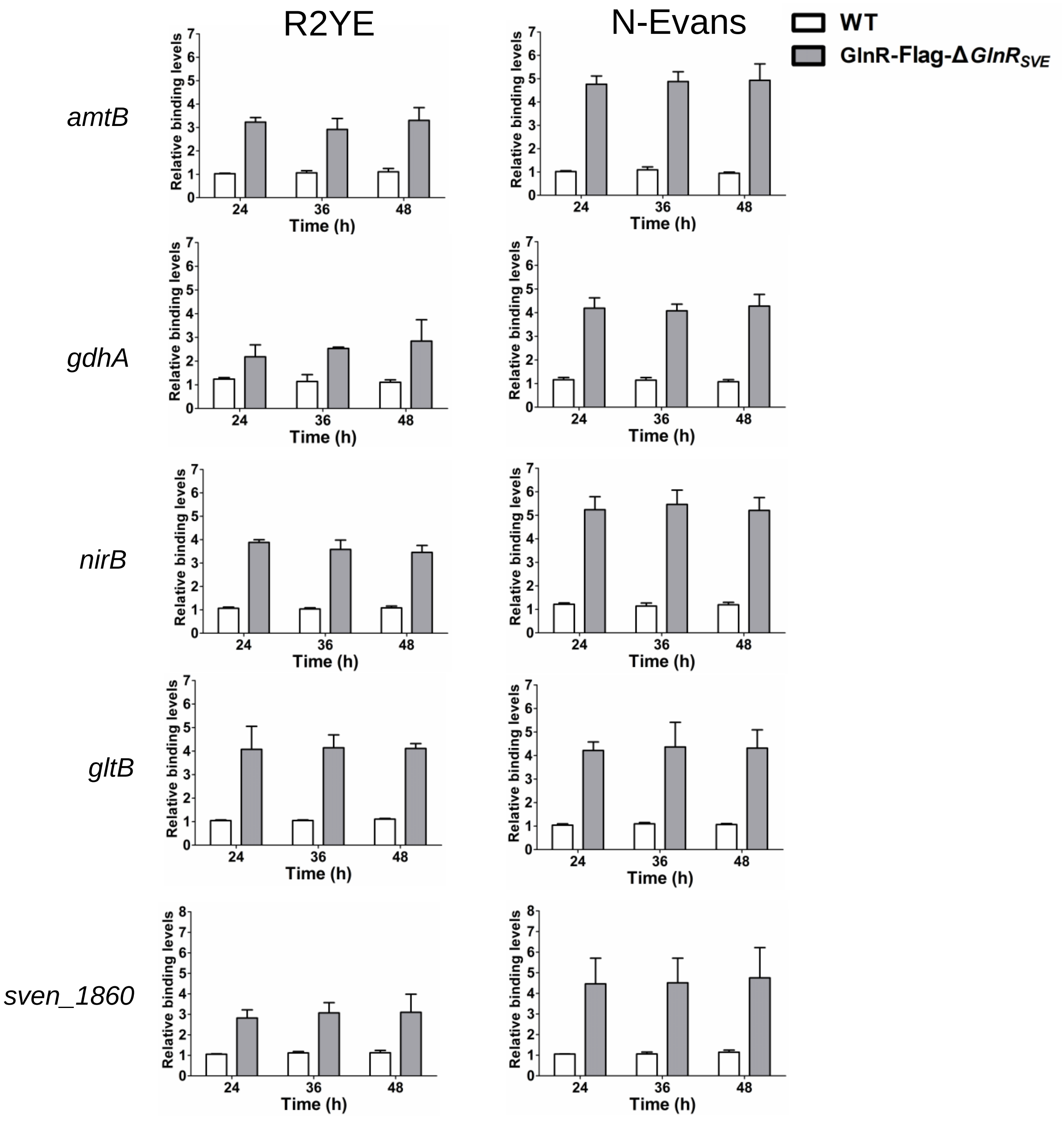


Figure S20. ChIP-qPCR analysis of the binding of GlnR to the promoters of nitrogen metabolism genes from cultures grown on R2YE or N-Evans. Analysis was performed using strains GlnR-FLAG-Δ$glnR_{SVE}$, and the wild-type 10712 (WT) grown on R2YE for N-Evans for the indicated times. The y-axis shows the binding levels of GlnR-FLAG in GlnR-FLAG-Δ$glnR_{SVE}$ and WT relative background levels, which was determined by recovery of target sequences. Results are the means (±SD) of triplet biological experiments.
